# Cognitive effects of placebo antipsychotics: Investigating their mechanisms using event-related potentials (ERPs) in a cognitive test and a self-referential task

**DOI:** 10.64898/2025.12.03.691789

**Authors:** Mingyi Diao, Ilya Demchenko, Jingyan Quan, Aidan Schottler-Raymond, Samuel Paul Louis Veissière, Jiani Zhang, J. Bruno Debruille

**Affiliations:** Douglas Mental Health University Institute, Montreal, QC H4H 1R3, Canada; Department of Neurosciences, McGill University, Montreal, QC H3A 2B4, Canada; Department of Psychology, McGill University, Montreal, QC H3A 1G1, Canada; Department of Psychology, Université du Québec à Montréal; Department of Anthropology, McGill University, Montreal, QC H3A 1A1, Canada; Department of Cognitive Science, McGill University, Montreal, QC H3A 1A1, Canada; Department of Psychiatry, McGill University, Montreal, QC H3A 1A1, Canada; Department of Physiology, McGill University, Montreal, Qc H3G 1Y6, Canada

**Keywords:** Placebo effects, Antipsychotics, Event-related brain potentials (ERPs), Cognition, Stimulus-self binding

## Abstract

Placebos modulate neurocognitive processes. We examined whether a fully deceptive antipsychotic-placebo changes cognition not only by enhancing arousal and attention but also by changing self-representations, namely, by adding that of being under a treatment related to psychosis and by weakening the binding of experience to reality. Drug-naive healthy participants (N = 83) were split into a group (N = 40) who took a pill described as an antipsychotic and a no-pill control group (N = 43). Participants performed a cognitive test (CogTest), that is, a semantic categorization task and a self-referential role judgment task (SRT). Behavioral responses and event-related potentials (ERPs) were recorded.

In the CogTest, placebos showed faster response times and higher accuracy than no-pills. This cognitive improvement was associated with more negative occipito-temporal early ERPs (OTEEs), larger central P2s (CentP2s), and larger late positive potentials (LPPs), with the latter correlating with response times. As in the literature, CentP2s were maximal in the SRT, possibly because it maximally stimulates the binding of the stimulus to long-term self-representations. The larger CentP2s and LPPs for placebos than for no-pills in the CogTest were thus tentatively related to the binding of the stimulus with egocentric self-representations, temporarily enriched by the addition of the representation of being on drug. On the contrary, the more negative OTEEs were related to a weaker binding to allocentric ones, preventing a strong binding of experience with reality. The smaller N400s observed for placebos than for no-pills in both tasks were related to greater openness to new experiences.

## 1. Introduction

Schizophrenia is characterized not only by positive and negative symptoms but also by cognitive deficits, particularly in semantic-, working-, and episodic-memory (Heinrichs & Zakzanis, 1998; Lee & Park, 2005; Yücel et al., 2012). These impairments are critical for long-term functional outcomes, as they impact social, occupational and daily life functioning (Allott et al., 2011; Bowie & Harvey, 2006; Green et al., 2000).

Coined by Bleuler in 1908, the term schizophrenia, literally meaning “split mind”, reflects a profound alteration in self-representations (Ashok et al., 2012; Bleuler, 1911; Hamm et al., 2017; Jeannerod, 2009; Lysaker & Lysaker, 2010; Sass & Parnas, 2003). Indeed, schizophrenia patients often feel themselves as being different (e.g., “mad” or “sick”). For example, patients may describe feelings of inhabiting another person’s body or a loss of boundaries between the self and the external world (Laing, 1994; Nordgaard et al., 2018). Individuals may feel possessed by foreign entities or invaded by the thoughts and voices of others (Hacker et al., 2008; Parnas & Handest, 2003).

Following antipsychotic treatments, the alleviation of such symptoms may be accompanied by cognitive improvements (Hill et al., 2010; Meltzer & McGurk, 1999). However, most antipsychotic medications primarily act by blocking dopamine receptors in the whole brain. Given that this disrupts many delicate neurochemical balances necessary for various cognitive processes, these medications appear unlikely to improve cognitive performance *directly* (Cools & D’Esposito, 2011; Kaar et al., 2020; Kapur & Seeman, 2001). The cognitive improvements observed alongside the symptom reduction may thus be due, at least in part, to their placebo effects (Hird et al., 2023; Kemp et al., 2010; Leucht et al., 2017).

However, such effects can be difficult to control for. Given their nature, they may vary according to who gives the pill. Prestige, such as the one of scientists working in renowned mental health research institutions affiliated with a university, may amplify the placebo effects merely induced by the idea that one can “get better” from taking pills. Moreover, in the case of psychoactive drugs, these effects might be boosted by what has been called “neuroenchantment” (Ali et al., 2014), a naive belief in the therapeutic power of neuroscience technologies (Kaptchuk, 2002; Olson et al., 2021; Olson et al., 2020). Taking a pill presented as a medication that changes the way the brain works in this highly suggestive “neuroenchanting” context may thrill participants and increase their arousal. The attention increase that may result could, in itself, be responsible for cognitive improvements.

Nevertheless, the effects of such an attention increase can be partially monitored by recording the reaction times (RTs) and the event-related brain potentials (ERPs) elicited by the stimuli of the cognitive tests used. Indeed, increased attention is typically associated with faster RTs and larger amplitudes of occipital P1s, N1s, and P2s in the case of visual stimuli (Carretié et al., 2004; Gupta et al., 2019; Luck, 1995, 2012) and larger N400s and late positive potentials (LPPs) regardless of the sensory modality of the stimuli used (Donchin & Coles, 1988; McCarthy & Nobre, 1993; Polich, 2007; Sergent et al., 2005).

On the other hand, placebos presented as psychoactive drugs can modify self-models by introducing the representation of being “under treatment.” This self-representational update is not merely semantic or abstract: it is encoded as a temporary shift in the embodied predictive model of the self that the brain uses to regulate perception, cognition, and action. From the standpoint of active inference (Ramstead et al., 2023), the brain continuously generates predictions about incoming sensory and interoceptive data based on its model of the world and of the self. When a placebo is administered in a context that suggests cognitive alteration (e.g., an antipsychotic), the brain integrates this input and updates its prior beliefs to include the notion that “I am now medicated by a drug that is related to psychosis and that impacts the way my brain works.” This temporary belief may thus influence attention, response selection, and stimulus interpretation.

The cognitive consequences of this self-update are realized through the dynamic binding of sensory stimuli to the updated self-model. When we perceive a stimulus, we are not simply observing it. We are also aware that it is *we* who are perceiving it. A moment in which the self is the agent experiencing the stimulus under certain internal states is constructed. In neurocognitive terms, this also means that representations activated by the stimulus are bound to representations of the self. This could explain why self-relevant stimuli (such as one’s own name or roles) evoke larger central P2s and late positive potentials (LPPs) (Fields & Kuperberg, 2012; Zhao et al., 2021). The placebo-induced addition to the usual self of being under the influence of a drug can amplify this binding process. In terms of ERP signatures, this could be reflected in larger central P2 and LPP ERPs, whereas a generalized arousal would imply not only these enlargements, but also those of the ERPs indexing the processing of the physical features of the stimulus, such as the occipital P1s and N1s for visual stimuli (Di Russo et al., 2002; Schendan et al., 1998) and of the central N1s for auditory ones (Lightfoot, 2016; Woods, 1995). Research has also highlighted the importance of the self in high-level cognitive functions, such as perception and attention (Northoff, 2016; Sui et al., 2013; Woźniak & Knoblich, 2019). Moreover, neuroimaging studies have shown that placebo effects are modulated by brain regions involved in top-down self-regulation processes that assign personal meaning to subjective experiences (Benedetti et al., 2005; Northoff et al., 2006; Schneider, 2007). Given that the mechanisms underlying placebo effects remain incompletely understood, Schneider and Kuhl (2012) wrote: “One promising way to approach placebo effects stems from a psychological analysis of the self conceived as a functional system.”

These perspectives complement and deepen the traditional expectancy view of placebo effects by highlighting how changes in self-representations modulate the contents of conscious experience (Benedetti, 2021). It is not just that one expects to perform better—it is that the self who performs is changed to some extent. The improved cognitive performance observed in the placebo condition may result from this enriched and updated self-model, rather than solely from increased attention or task engagement.

The present study thus investigated whether positive cognitive effects of placebos could also result from the changes in the current representations of the self that are associated with the belief of being under the influence of a drug. To achieve this goal, drug-naive healthy participants were recruited to eliminate both expectations of prior interoceptive experiences of the drug and the hope of symptom relief. The pill was presented as an antipsychotic medication, as being either risperidone or olanzapine and described as a tranquilizer to decrease expectations of cognitive enhancement. A separate survey confirmed that the informed consent form used in this study effectively mitigated non-specific placebo expectations (see Text S9-S10 & Table S6-S10 of Supplementary Materials). RTs and ERPs were recorded in a cognitive test and in a self-referential task to assess changes in self-representations and to control for arousal.

## 2. Methods and Materials

### 2.1 Participants

Eighty-three right-handed anglophone participants were recruited through online advertisements (e.g., Kijiji, Facebook and the McGill Classified Ads website). The placebo (*N* = 40) and the no-pill groups (*N* = 43) were recruited in separate cohorts. The advertisement for the placebo group explicitly stated that upon their arrival to the lab, participants would take a single minimal dose of an antipsychotic. All participants had at least twelve years of education in English and normal or corrected-to-normal vision. They reported having no psychiatric history (except for a depressive episode that resolved at least two years ago), no neurological condition compromising brain functioning, no intellectual deficits, and no current use of psychiatric medication. All participants were given one of the two informed consent forms prior to their participation, and the data were processed anonymously. The informed consent forms for no-pill participants as well as those for placebo participants (see Text S1–S2 of Supplementary Materials) were approved by the Douglas Ethics Review Board (project number: IUSMD-06-42). All research was performed in accordance with relevant guidelines/regulations and in accordance with the Declaration of Helsinki.

### 2.2 Psychometric scales

Before the testing, participants filled out a demographic form. The evolution of their anxiety and fatigue was measured by using the state portion of the state-trait anxiety inventory form Y (the STAI-Y State) (Spielberger, 1983) and a lab-made fatigue questionnaire (see Text S3-S5 and Table S1-S3 of Supplementary Materials). The schizotypal personality questionnaire (SPQ) was used to assess their schizophrenia attributes. The STAI-Y has high reliability coefficients, 0.92 for the state- and 0.90 for the trait-scale (Form Y), respectively. The STAI-Y and the fatigue questionnaires had to be completed twice, that is, once before and once after the experiment. The SPQ, which is based on the symptoms of schizophrenia, was initially designed to measure the severity of schizotypal personality traits in the general population (Raine, 1991). It is a self-reported questionnaire that contains seventy-four items with high internal reliability (alpha > 0.90) and test-retest reliability (r = 0.82). It was administered before the experiment and only once since SPQ total scores remain relatively stable over time (Moreno-Izco et al., 2015).

### 2.3 Stimuli

Each participant was asked to complete the two tasks. The cognitive test was the semantic categorization used by Prévost et al. (2011) and Diao et al. (2024). It was selected because of its simplicity and because the behavioral and neurocognitive indexes it produces cover vigilance, visual attention, processing of visual features of stimuli, semantic memory, working memory, automatic encoding in episodic memory and decision-making processes (Debruille et al., 2007; Diao et al., 2024), which can all be improved by attention increases. This task employs linguistic stimuli and focuses on understanding their meaning, as language comprehension is key to patients’ psychosocial rehabilitation. This task was also chosen because it reveals behavioral and neurocognitive deficits in people with schizophrenia attributes (SzAs, i.e., schizophrenia patients and subclinical people with schizotypal traits) (Debruille et al., 2007; Debruille et al., 2013; Prévost et al., 2011). It includes 180 trials in total, each made of two serially presented words. Two-thirds of these trials start with the question word “ANIMAL?” presented in the center of the screen for 500 ms. It is replaced by a fixation cross for 500 ms. Then, an exemplar (e.g., dog) or a non-exemplar (e.g., table) target word appears for 1000 ms (these stimuli and the timing of their presentations are in Table S12 and Figure S1 of Supplementary Materials, respectively). Participants are asked to decide whether or not the target word belonged to the animal category as accurately and fast as possible by pressing with their right index finger a “Yes” button for matching targets or a “No” button for mismatching targets. Each target word is then replaced by a blank screen for 500 to 1000 ms. After that, the word “Blink” is presented and lasts for 500 ms. Exemplar and non-exemplar words are matched for the mean number of letters and frequency of usage.

The second task was the social role acceptance task used by Fernandez et al. (2016) and Diao et al. (2023). In this task, previous studies showed that participants with subclinical schizotypy do not show the poorer neurocognitive indexes (e.g., slower response times) that are typically observed in cognitive tests in such participants with schizophrenia attributes (Diao et al., 2023; Fernandez-Cruz et al., 2016). This lack of deficits could be due to the nature of this task, which involves self-related choices and extraordinary social roles. It allows individuals with high schizotypy to express the variety and peculiarities of their social drives. It probably also increases attention by being more attractive and motivating for them than classical cognitive tests. Here, the reason for choosing this social role task was also its self-referential aspect, which maximally deepens the association of the stimulus with the self. This contrasts with the minimum binding of the stimulus to the self that automatically occurs in usual cognitive tests. There, people are consciously perceiving the stimulus and simply aware, at the same time, that it is they who are perceiving it (Baars et al., 2003; Gallagher, 2000; Merleau-Ponty et al., 2013). The larger P2 ERPs found in self-referential than in non-self-referential tasks (Xu et al., 2017; Zhao et al., 2021) could index this deeper association of stimuli with the self. Indeed, as mentioned, these larger P2s may not be due to a mere increase of attention as they may not be associated to larger amplitudes of the ERPs indexing the processing of the physical features of the stimuli, such as the occipital P1 and N1s for visual stimuli, whereas these amplitude enlargements are seen when attention to those stimuli is increased. Two subsets of 200 names of social roles were extracted from the original set of 401 names used in Fernandez et al. (2016). Each participant was presented with only one of these subsets. Each trial was made of one social role name presented for 500 ms at least, and 1800 ms at most (these stimuli and the timing of their presentations are in Table S13 and Figure S2 of Supplementary Materials, respectively). Participants were asked to decide as quickly as possible whether or not they would consider performing the role at any moment in their life by pressing a “YES” or a “NO” button using their right index and middle finger, respectively. The name was replaced either by a fixation cross or by a “BLINK!” stimulus.

### 2.4 Experimental Design

Upon arrival, participants were informed of the purpose of the study and signed an informed consent form. For the placebo group, this form stated that participants would receive an antipsychotic medication, namely, 1 mg of risperidone or 2.5 mg of olanzapine. The possible therapeutic and side effects associated with this medication were described. The verbatim text for the therapeutic effects was: “It is known that antipsychotic medications function to improve a number of psychotic symptoms. The exact mechanisms of this function remain incompletely understood. One possibility is that these medications facilitate, directly or indirectly, the mechanisms that allow us to understand unexpected information. By doing so, these medications could help patients with inaccurate beliefs (e.g., delusions) to change their minds.” The verbatim text for the side effects was: “Risperidone (or olanzapine) is a drug that is widely used in clinical practice by thousands of patients and is approved by the Food and Drug Administration. The (1 or 2.5) milligram dose you will be taking is almost the lowest dose that is given to adults. There are several possible side effects associated with this medication, although it is extremely unlikely that they could occur at the dose involved in this study. Also, note that all studies that examined side effects looked at repeated administrations of the drug, rather than at a single dose. The adverse effects of this dose of risperidone or olanzapine that you might experience are somnolence, dry mouth, light-headedness, constipation, increased appetite, stomach upset (nausea/indigestion), restlessness, sense of muscle weakness (with no actual loss of strength), insomnia, and muscle stiffness. However, mild drowsiness is the only adverse effect that is likely to occur.”

Subsequently, participants completed a set of demographic and psychometric scales. The electrode cap was then placed. Participants were seated comfortably in a dimly lit room, 70 to 100 cm from the computer screen. After task instructions (see Text S6-S7 of Supplementary Materials) and a short practice session, the placebo group received a pill that looked like a risperidone or olanzapine tablet but contained saccharose. The participants of the no-pill group did not receive any tablets. Immediately after, the RT and EEG recording session, which included the cognitive test and the social role task, began.

### 2.5 Data acquisition

As mentioned, reaction times (RTs) were recorded in both tasks. Reaction accuracies (RAs) were recorded in the cognitive test, whereas the nature of responses (i.e., role acceptance vs. rejection) was recorded in the social role task. The electroencephalogram (EEG) was captured with an Electro-Cap International cap including 28 tin electrodes placed at the Fp1/2, F3/4, Fc3/4, C3/4, Cp3/4, P3/4, O1/2, Fz, Fcz, Cz, Pz, F7/8, Ft7/8, T3/4, Tp7/8, and T5/6 locations of the international 10-20 system. The right earlobe was used as a reference and the ground was placed 2 cm anterior to the Fz site. Impedances were kept below 5 KΩ. The 60 Hz EM noise coming from the power lines was reduced by an electronic notch filter. High- and low-pass filters had their half amplitude cut-off set at 0.01 Hz and 100 Hz, respectively. Signals were digitized at a 248 Hz frequency.

### 2.6 Data processing and measures

Electrophysiological data were processed with the EEGLAB and the ERPLAB toolbox for MATLAB. Processing and study-specific details are included in prior works (Diao et al., 2024; Diao et al., 2023) and in Text S8 of Supplementary Materials of the present study. ERPs at each electrode were calculated by averaging all the accepted EEG epochs of each condition (mismatch vs. match or rejection vs. acceptance). Only participants with at least 30 accepted trials in each condition were retained. Table S11 of Supplementary Materials provides, for the remaining 83 participants, the average number of accepted trials.

The amplitudes of the P2, N400, and late positive potential (LPP) were measured, as they are the most cognitive potentials, that is, those that occur right after the potentials that depend in a large part on the sensory modality (e.g., visual or auditory) of the stimulus. The P2 depends on the nature of the stimulus (e.g., the P2 elicited by a written word differs from that evoked by a picture of a face). Its amplitude is larger when more attention is allocated to the processing of the stimulus (Carretié et al., 2004; Gupta et al., 2019) and, most importantly, when this stimulus occurs in a self-referential task (Fields & Kuperberg, 2012; Zhao et al., 2021). On the other hand, the amplitude of the N400 is well-known to index the difficulty of semantic processing (Kutas & Federmeier, 2011; Renoult et al., 2012). This amplitude also depends on the strategy of processing that develops over the task (Chwilla et al., 1995). Finally, the amplitude of the LPP is positively correlated with attention and with the amount of information placed in working memory (Donchin & Coles, 1988; Polich, 2007; Sergent et al., 2005) and, thus, in consciousness (Hartigan & Richards, 2017; Kutas et al., 1977). As such, it is also larger in self-than in non-self-referential tasks (Xu et al., 2017; Zhao et al., 2021). To account for differences in attentional resources allocated to the processing of the visual features of the stimuli, the amplitudes of the occipito-temporal N1s were also measured. All the amplitude measures consisted of the mean voltages of the ERPs during specific time windows: 145 ms to 270 ms post-stimulus for the occipito-temporal N1 at T5/6 and O1/2 electrodes, 180 to 270 ms for the P2 at all electrodes except T5/6 and O1/2, 270 to 500 ms for the N400 at all electrodes, and 500 to 800 ms for the LPP at all electrodes. Table S11 of Supplementary Materials provides the mean amplitudes of the N1, P2, N400, and LPP in each condition.

### 2.7 Statistical analysis

Mixed-model repeated-measure ANOVAs were run to analyze the RTs, RAs and ERPs with group (no-pill vs. placebo) as the between-subject factor. The within-subject factors for the two tasks were the conditions (mismatch vs. match for the cognitive test and rejection vs. acceptance for the social role task) and the electrodes for ERPs. As previous literature (Xu et al., 2017; Zhao et al., 2021) has shown that P2s and LPPs are larger in self-referential-than in non-self-referential-tasks, we compared these two ERPs and RTs using task (cognitive test vs. social role task) and condition (mismatch/rejection vs. match/acceptance) as within-subject factors. Pearson’s r correlation analyses were used to detect associations between RTs and ERP measures in both tasks and both groups. One-tailed tests were used given the *a priori* hypothesis that larger ERP positivities correlate with faster RTs, as previously found (Bamford et al., 2015; Xing et al., 2022). All analyses were done with IBM SPSS Statistics (version 27). Degrees of freedom were adjusted with Greenhouse and Geisser’s procedure to compensate for the heterogeneity of variances across electrodes. In those cases, the original F values and degrees of freedom are provided together with the corrected p values. Effect sizes were reported as the proportion of variance explained by the variable (*ηp^2^*). The Benjamini-Hochberg false discovery rate (B-H FDR) procedure was used to judge the statistical significance of the p-values of each series of tests. The false discovery rate chosen was 10%. All figures were created using GraphPad Prism 9 and refined with Inkscape.

## 3. Results

### 3.1 Demographics

The demographics of the participants are shown in Table 1. The no-pill and placebo groups did not differ significantly in terms of sex ratio, mean age, and mean number of years of education. There were significant interactions of time x group on the STAI anxiety level (F (1, 81) = 25.1, *p* = 3.2 x 10^-6^, *ηp^2^* = 0.24) and on the fatigue level (F (1, 80) = 15.1, *p* = 2.1 x 10^-4^, *ηp^2^* = 0.16). Anxiety of all participants significantly increased along with the experiment irrespective of the group assignment, but this increase was larger for the no-pill than for the placebo group. As to the fatigue level, it significantly increased along with the experiment only in the placebo group. After the experiment, the mean fatigue level of this group was significantly higher than that of the no-pill group.

**Table 1:** Demographic and clinical characteristics of participants. ** are for p < 0.01. *** are for p < 0.001.

|  |  | <u>No-Pill</u><br>(*N* = 43) | <u>Placebo</u><br>(*N* = 40) |
| --- | --- | --- | --- |
| Biological Sex: male | % (*N*) | 46.5% (20.0) | 47.5% (19.0) |
| Age | *M* (*SD*) | 23.0 (3.1) | 22.8 (3.1) |
|  | Range | 18-30 | 18-30 |
| Race | Asian (*N*) | 20 | 9 |
|  | White (*N*) | 11 | 22 |
|  | Black (*N*) | 1 | 3 |
|  | Hispanic (*N*) | 5 | 1 |
|  | Mixed (*N*) | 3 | 2 |
|  | Middle eastern (*N*) | 3 | 1 |
|  | Missing (*N*) | 0 | 2 |
| Years of Education | *M* (*SD*) | 15.1 (1.7) | 14.9 (1.6) |
|  | Range | 11.0-18.0 | 12.0-18.0 |
| Total SPQ | *M* (*SD*) | 17.9 (11.1) | 18.9 (15.2) |
|  | Range | 0-38 | 0-58 |
| Anxiety (pre-experiment) | *M* (*SD*) | 22.4 (17.4) | 22.7 (14.6) |
|  | Range | 0-64.3 | 0-61.9 |
| Anxiety (post-experiment) | *M* (*SD*) | 57.7 (5.2) *** | 35.1 (24.2) |
|  | Range | 47.6-66.7 | 0-95.2 |
| Fatigue (pre-experiment) | *M* (*SD*) | 31.0 (19.6) | 29.9 (16.4) |
|  | Range | 0-75.0 | 0-66.7 |
| Fatigue (post-experiment) | *M* (*SD*) | 30.7 (19.2) ** | 44.8 (24.9) |
|  | Range | 0-75.0 | 0-100.0 |

### 3.2 Cognitive test (Semantic categorization task)

#### 3.2.1 Behavioral results

RAs of the placebo group (Mean = 95.8%, SD = 3.6) were slightly but significantly higher (F (1, 81) = 6.9, *p* = 0.01, *ηp^2^* = 0.078) than those of the no-pill group (Mean = 93.1%, SD = 8.1). RTs of the placebo group (Mean = 751 ms, SD = 112) were notably faster (F (1, 81) = 13.6, *p* = 4.0 x 10^-4^, *ηp^2^* = 0.14) than those of the no-pill group (Mean = 826 ms, SD = 86). RTs were shorter in the match-(Mean = 766 ms, SD = 102) than in the mismatch-condition (Mean = 814 ms, SD = 104) (F (1, 81) = 63.7, *p* = 8.2 x 10^-12^, *ηp^2^* = 0.44) (Figure 1).

**Fig. 1.**
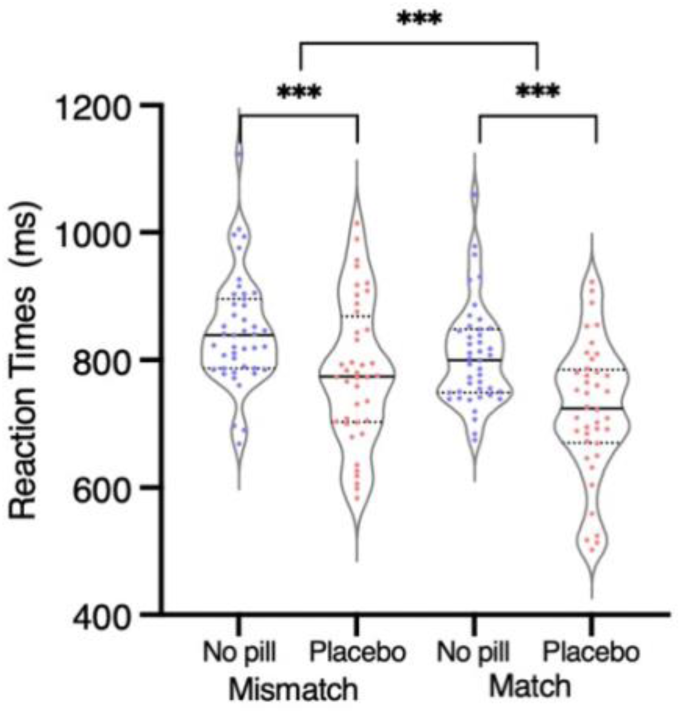
Mean reaction times for the mismatch- and the match-condition obtained in the cognitive test (that is, the semantic categorization) for each of the participants of the no-pill group (N = 43) and for each of those of the placebo group (N = 40). The solid black lines are medians. The thin dotted black lines are quartiles. *** are for p < 0.001.

#### 3.2.2 Electrophysiological results

##### 3.2.2.1 N1 amplitudes

The ANOVA revealed a significant group x electrode interaction (F (3, 243) = 3.7, *p* = 0.023, *ηp^2^* = 0.043). Placebo participants (Mean = -1.0, SD = 2.4) had larger N1 amplitudes than no-pill participants at T5/6 and O1/2 (Mean = 0.39, SD = 2.1) (see Table S4 of Supplementary Materials).

##### 3.2.2.2 P2 amplitudes

The ANOVA revealed a significant electrode x group interaction (F (23, 1863) = 6.0, *p* = 2.4 x 10^-5^, *ηp^2^* = 0.069). The post hoc ANOVAs run at each electrode to find the source of this interaction indicated that placebo participants had larger P2 amplitudes at Fp2, F8, Fz, Cz, F4, Fc3 and Fcz than no-pill participants (Figure 2). The opposite result was found at Tp8 (see Figure S2 below and Table S4 of Supplementary Materials for p-values for each electrode).

**Fig. 2.**
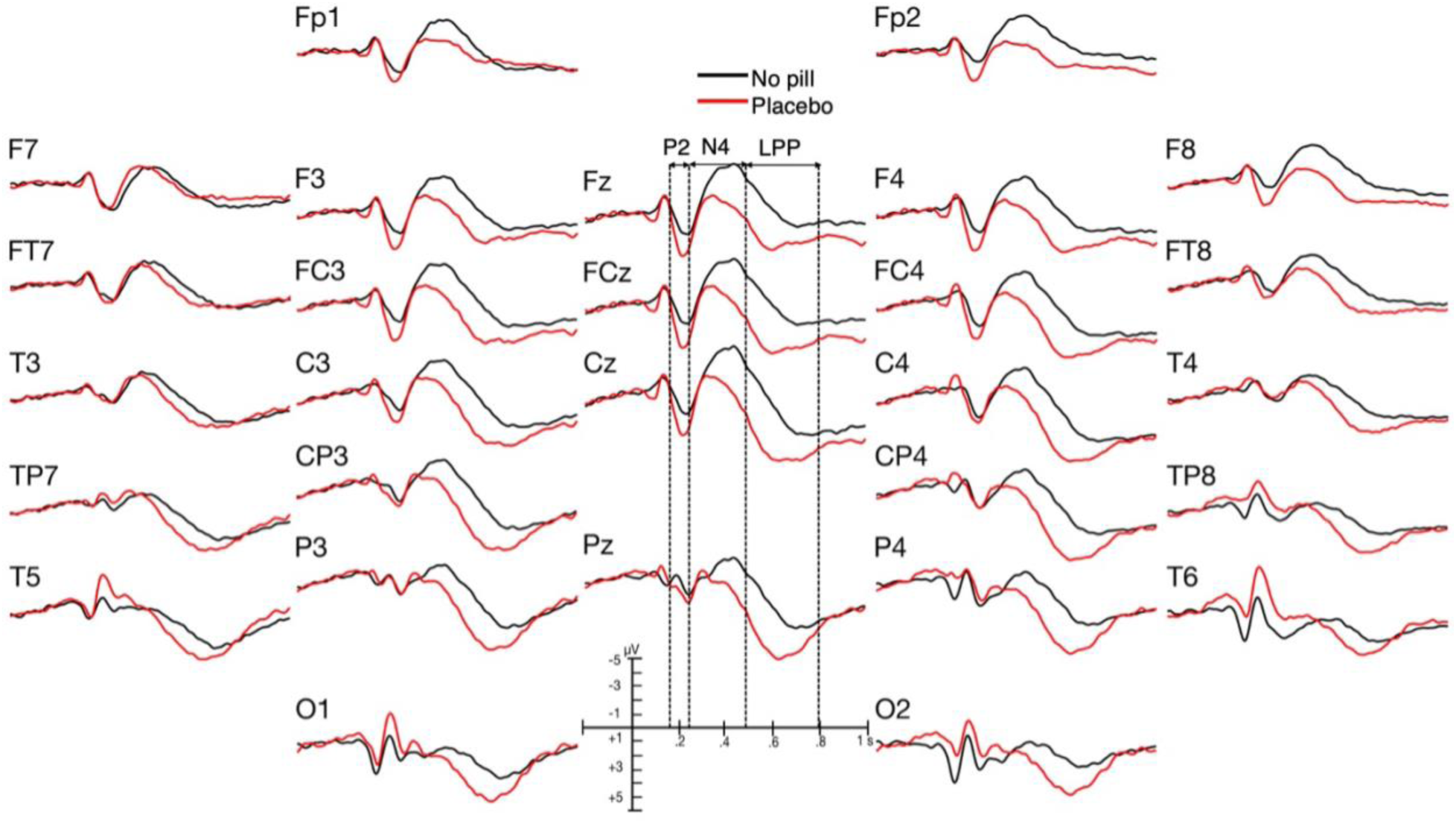
Grand average ERPs of the placebo group (N = 40, red lines) and of the no-pill group (N = 43, black lines) in the cognitive test.

##### 3.2.2.3 N400 amplitudes

The analysis revealed that group interacted with electrode (F (27, 2187) = 4.0, *p* = 0.003, *ηp^2^* = 0.047). Participants who took the placebo had significantly smaller (i.e., less negative) N400 amplitudes at Fp2, F8, Fz, Cz, Pz, F4/3, Ft8, Fc4/3 and Fcz compared to those who did not take any pill (Figures 2 and 3). N400 amplitudes of the mismatch condition (Mean = -1.2 µV, SD = 2.8) were significantly larger (F (1, 81) = 47.4, *p* = 1.1 x 10^-9^, *ηp^2^* = 0.37) than those of the match condition (Mean = -0.21 µV, SD = 3.0).

**Fig. 3.**
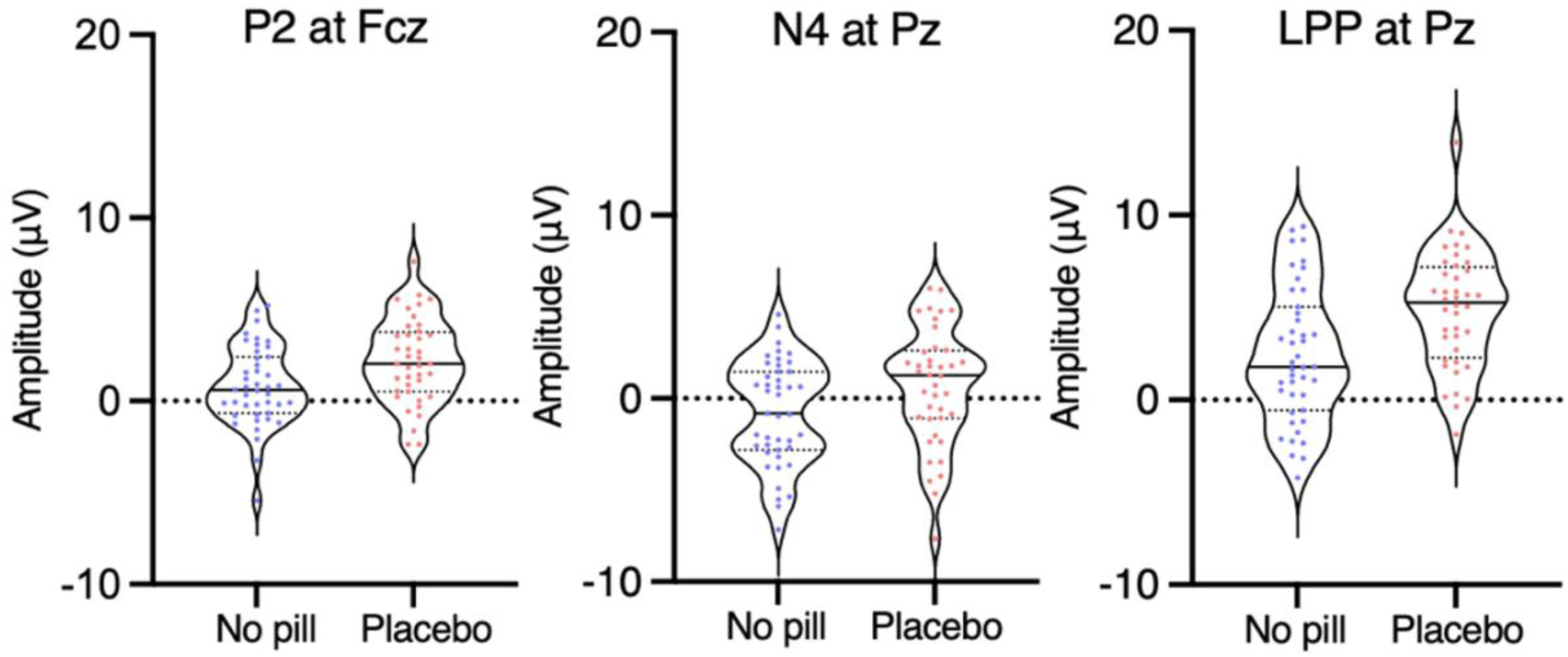
Amplitudes of P2s at Fcz, of N400s at Pz and of LPPs at Pz in the cognitive test for each of the participants of the no-pill group (N = 43, blue plot) and for each of those of the placebo group (N = 40, red plot). The solid black lines are medians. The thin dotted black lines are quartiles.

##### 3.2.2.4 LPP amplitudes

The ANOVA revealed interactions of electrode with group (F (27, 2187) = 3.4, *p* = 0.007, *ηp^2^* = 0.04) and of semantic condition with group (F (1, 81) = 6.3, *p* = 0.014, *ηp^2^* = 0.072). The post hoc ANOVAs revealed that LPP amplitudes of the placebo group were significantly larger than those of the no-pill group at Fp2, F8, Fz, Cz, Pz, P4/3, T6/5/4, F4/3, Ft8, Fc4/3, Fcz, C4/3, Tp8/7, Cp4/3 and O2/1 in the mismatch condition, and at Fp2, F8, Cz, Pz, P4, Ft8, Fc4, Fcz, C4/3, Tp8, Cp4/3 and O2 in the match condition (see Table S4 of Supplementary Materials).

#### 3.2.3 Correlations between RTs and ERPs

In the placebo group, there was no strong correlation between RTs and P2 amplitudes, that is, no correlation coefficient that was larger than |0.4|. Stronger correlations were found between RTs and N400 amplitudes at Fz (Pearson’s r = -0.65, *p* = 2.6 x 10^-6^), Fcz (Pearson’s r = -0.63, *p* = 7.0 x 10^-6^) and Cz (Pearson’s r = -0.58, *p* = 4.0 x 10^-5^) in the match condition. A similar pattern was observed in LPP amplitudes where RTs were significantly correlated with LPP amplitudes in both match and mismatch conditions at Cz (Pearson’s r = -0.66, *p* = 1.6 x 10^-6^ and -0.56, *p* = 8.0 x 10^-5^, respectively), Fcz (r = -0.66, *p* = 1.5 x 10^-6^ and -0.57, *p* = 5.6 x 10^-5^, respectively), and Fz (-0.52, *p* = 3.2 x 10^-4^ and -0.54, *p* = 1.9 x 10^-4^, respectively), as illustrated by Figure 4. In the no-pill group, no strong correlation was found between RTs and the amplitudes of the three ERPs, that is, all correlation coefficients were smaller than |0.3|.

**Fig. 4.**
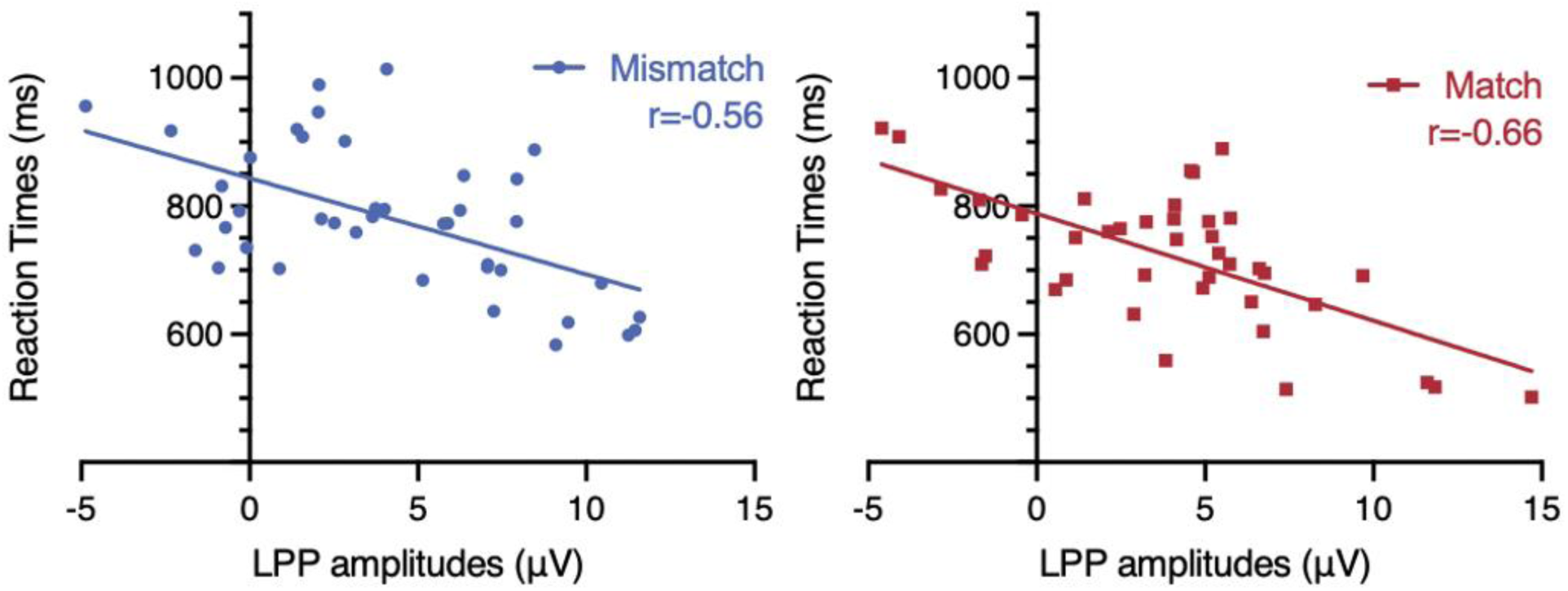
Correlations between reaction times and ERPs in the placebo group in mismatch and match conditions of the cognitive test. The x coordinate of each circle-participant is its mean LPP amplitude at Cz and the mean RT is its y coordinate (N = 40).

### 3.3 Social role self-referential task

For this task, there was no significant effect of group on RTs, N1-, P2- and LPP-amplitude (Figures 5 and 6). There was no strong correlation between RTs and ERP amplitudes in both groups, that is, no correlation coefficient was larger than |0.4|. Nevertheless, the ANOVA performed on N400 amplitudes revealed an effect of group (F (1, 81) = 8.2, *p* = 0.005, *ηp^2^* = 0.092). Placebos group had smaller N400 amplitudes (Mean = 0.41 µV, SD = 2.4) than no-pills (Mean = -0.64 µV, SD = 2.5). There was an effect of decision on N400 amplitudes (F (1, 81) = 8.0, *p* = 0.006, *ηp^2^* = 0.089). At all electrodes, they were found to be slightly larger for social role rejections (Mean = -0.27 µV, SD = 2.5) than for acceptances (Mean = -0.005 µV, SD = 2.5).

**Fig. 5.**
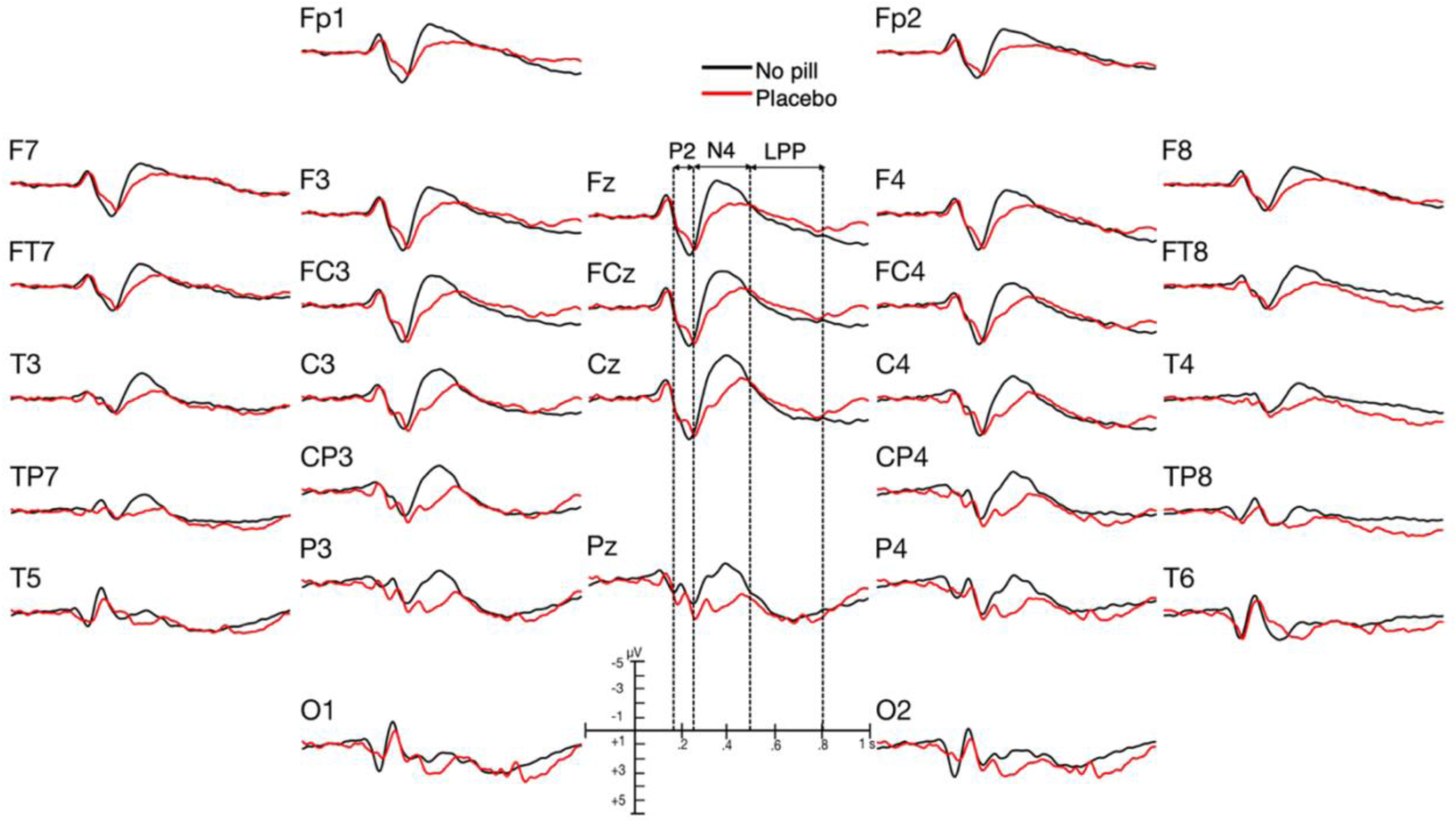
Grand average ERPs of the placebo group (N = 40, red lines) and of the no-pill group (N = 43, black lines) in the social role task.

**Fig. 6.**
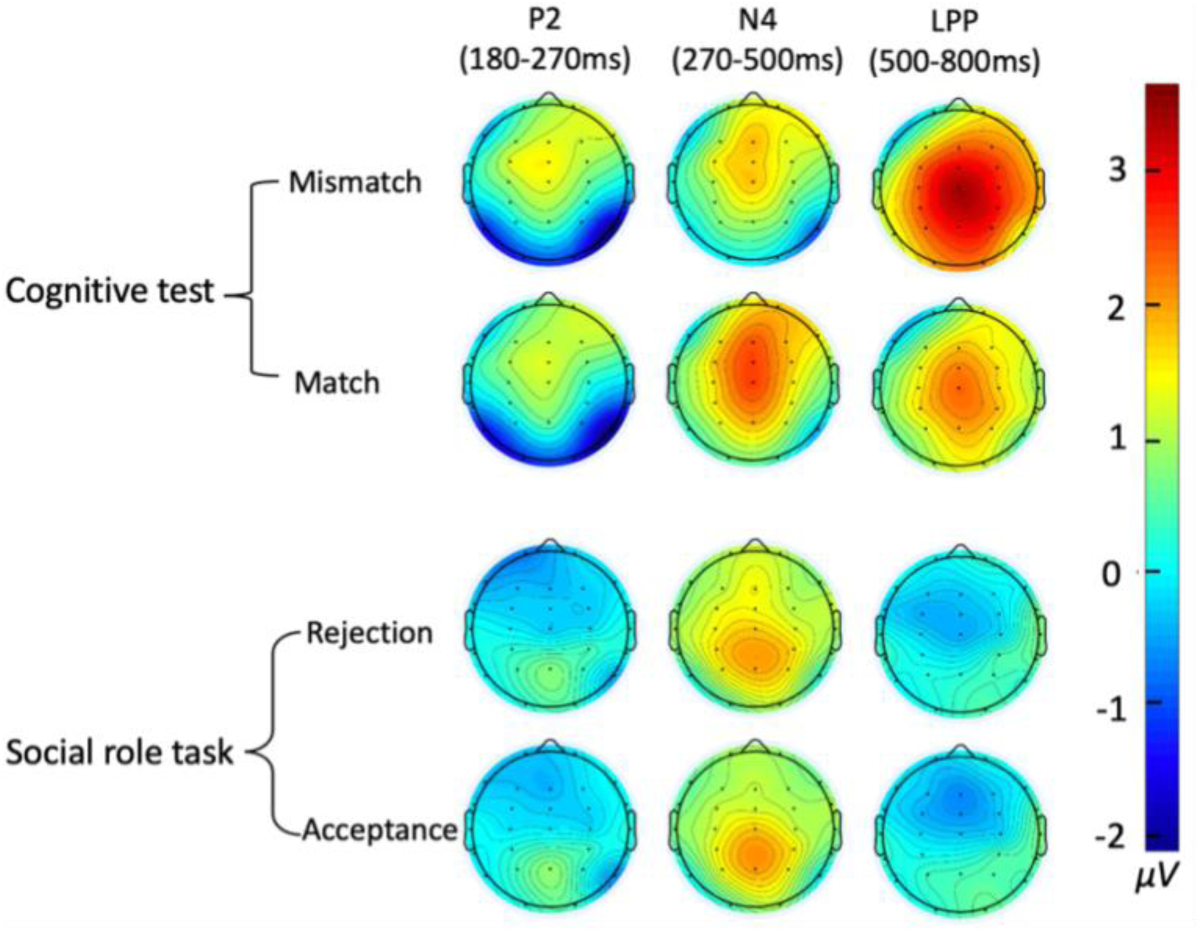
Spline interpolated iso-voltage scalp maps of the subtractions, in each time window, of the mean ERP voltages of the no-pill group (N = 43) from those of the placebo group (N = 40) in the cognitive test and the social-role self-referential task.

### 3.4 Comparison of the two tasks

RTs of the cognitive test (Mean = 790 ms, SD = 106) were much faster (F (1, 81) = 254.5, *p* = 1.0 x 10^-26^, *ηp^2^* = 0.76) than those of the social role task (Mean = 1017 ms, SD = 164). Participants had larger P2 amplitudes in the social role task than in the cognitive test at Fc3/4, C3/4, Cp3/4, P3/4, Fcz, Cz, Pz, F8, Ft7/8, T3/4 and Tp7/8 (task x electrode interaction, F (23, 1886) = 3.2, *p* = 0.015, *ηp^2^* = 0.037). There was also a task x condition x electrode interaction for LPP measures (F (27, 2214) = 3.2, *p* = 0.004, *ηp^2^* = 0.037). LPP amplitudes were larger in the cognitive test than in the social role task at Fp1, F7, Cz, Pz, P4/3, T6/5/4/3, F4/3, Ft7, Fc4/3, Fcz, C4/3, Tp8/7, Cp4/3 and O1 in the mismatch condition and at F7, Cz, Pz, P4/3, T6/5/4/3, F4, Ft8/7, Fc4/3, Fcz, C4/3, Tp8/7, Cp4/3 and O2/1 in the match condition (see Figure S3 of Supplementary Materials where the ERPs of these two tasks are superimposed, and Table S5 for p-values at each electrode).

## 4. Discussion

This study aimed to test if the intake of a placebo impacts cognition through changes of self-representations, such as the addition of the representations of being under the influence of a drug. To achieve that goal, we used a setting that, a) prevents hope of clinical improvements and expectations of an already felt interoceptive experience of the drug and, b) diminishes expectations of cognitive enhancements by presenting the pill as a tranquilizer. We recruited drug-naive healthy participants. We compared those who took a fully deceptive antipsychotic placebo to those who had no-pill. Two tasks were used: a cognitive one, namely, an animal versus object semantic categorization task and a particular social role self-referential task, where there was no correct or incorrect answer. We recorded response accuracies (RAs) in the cognitive test, response nature in the self-referential task, and response times (RTs) and event-related potentials (ERPs) in both tasks.

As in previous literature (Kutas & Federmeier, 2011; Xu et al., 2017), central P2s were larger in the self-referential task than in the cognitive test, and, in both tasks, N400 ERPs were found to be of smaller amplitude for the match/acceptance-than the mismatch/rejection-condition.

The placebo was found to improve cognition. In the cognitive test, RAs were a bit higher and RTs were notably faster than for no-pill participants. These improvements allow us to discard negative expectations that could have arisen from the prevalent attitudes toward antipsychotics (Helbling et al., 2006). These medications are seen as major tranquilizers and are often associated with slowed thought processes.

The examination of event-related potentials (ERPs) in the cognitive test revealed that placebos had larger occipital N1s, central P2s and late positive potentials (LPPs) than no-pills. At first glance, it seems that they could, together with the faster RTs, be accounted for by an attention increase related to the thrill generated by the neuroenchantment previously mentioned (Ali et al., 2014; Olson et al., 2021; Olson et al., 2020). This thrill could augment the effect of a placebo presented as a psychoactive drug. The informed consent form for placebo participants, which stated that the medication could “help patients with inaccurate beliefs”, may also have triggered positive expectations of more accurate thinking.

However, this attention interpretation cannot explain all the results observed. In the social role task, where both groups were exposed to the same neuroscientific laboratory context, neither the occipital P1 and N1, nor the central P2, or the LPP ERPs were enlarged by the placebo. These facts suggest that this task induced maximal visual attention in both groups, possibly because of the extraordinary stimuli it includes and the personal involvement it requires. In any case, it seems that if placebo participants were actually paying more visual attention than no-pill participants, placebos should also have had larger P1s than no-pills (Clark & Hillyard, 1996; Natale et al., 2006; Slagter et al., 2016; Taylor, 2002). This was not the case. On the contrary, occipital P1s appeared larger for no-pills than for placebos in both tasks (Figure S3 of Supplementary Materials).

Another possibility may then be discussed. These larger N1s could in fact be seen as less positive ERPs for placebos than for no-pills at temporo-occipital sites in the time-window of the central P2. They could be interpreted in the same theoretical framework as the more positive temporo-occipital ERPs recently found by Sinha et al. (2025) when participants are only with themselves. Namely, they could index variations of the strength of the binding of the stimulus with allocentric representations of the self (Chechlacz et al., 2010; Chen & Crawford, 2020; Chen et al., 2014; Neggers et al., 2006; Zaehle et al., 2007). Accordingly, the less positive ERPs found here could indicate a weaker binding of stimulus representations with this type of self-representations. This could make sense as it may be important to prevent a maximal binding of the currently activated representations with these allocentric representations of the self. Indeed, these representations should not fully include information that could be biased by the effect of the drug on the participant.

In contrast, the more positive central P2s observed in the cognitive test for placebos than for no-pills can be related to an increase binding with another type of self-representations, namely, with egocentric self-representations temporarily enriched by the addition of the representations of being on a drug. Indeed, larger central P2s are observed in tasks that can only increase the binding with such representations of the self, that is, in self-referential tasks (Chen et al., 2011; Fields & Kuperberg, 2012; Zhao et al., 2021). Like other automatic phenomena, such as the knee reflex, which can be increased or reduced, the automatic binding of stimulus representations with self-representations previously mentioned, is likely to be adaptable. The maximally large central P2s for both placebos and no-pills were observed in the social role task where the binding with the self can already be at the ceiling due to the self-referential nature of that task (Figure S3 of Supplementary Materials).

In addition to the central P2s, the LPPs were also larger in placebos than in no-pills. This was observed in the cognitive test and thus, when the binding was not at the ceiling. Like the larger central P2s, these larger LPPs may thus also be related to the binding of stimulus representations with a larger set of egocentric self-representations. However, instead of involving representations of the stimulus itself, as for the central P2s, the binding with the self should, for the LPP, pertain to representations corresponding to the meaning of this stimulus for the task at hand. Indeed, the LPP is thought to index the conscious evaluation of the stimulus in the task, namely, the representations of its meaning in its context of occurrence (Hartigan & Richards, 2017; Sergent et al., 2005). This is supported by the correlations found between the latency of the LPP peak and RTs when participants have to prioritize the accuracy of their response (Kutas et al., 1977). This is also supported by the existence of the binding of representations that correspond to the meaning of the stimulus for the task at hand with self-representations. Indeed, when people are conscious of this meaning, they are also conscious that it is they who are aware of it. Enriching the egocentric self might thus also increase the amplitudes of the LPPs evoked by the stimuli. Conversely, impoverishing it to the representation of an avatar in a virtual reality diminishes it (Spanlang et al., 2018).

The faster RTs observed in the cognitive test for placebos than for no-pills could be observed because such a wider self-binding with egocentric self-representations can only benefit the global level of activation of the representations of the stimulus, including those of the actions associated to it. This is revealed by the faster response times that occur when the various features of a stimulus are bound in a coherent whole in contrast to partial bindings and fragmented stimulus processing (Hasson et al., 2008; Mayerhofer & Schacht, 2015). Here, to explore this possibility, RTs were correlated with the amplitude of LPPs. The larger the mean amplitude of the LPP of a participant in the placebo group, the faster his/her mean RT was in the cognitive test, which aligns with previous studies (Bamford et al., 2015; Xing et al., 2022). This positive correlation was particularly strong at fronto-central sites (Fcz), in accordance with embodied views of cognition (Ansorge et al., 2010; Clark, 2008; Foglia & Wilson, 2013), which specify that the meaning of the stimulus is also coded by its affordances (Brakus, 2008; Shin, 2017; Soylu, 2016). These RT-ERP correlations become notable and significant in the placebo participants, which could be due to their larger LPPs, and which can be related to an increase in activation of stimulus representations induced by the wider self-binding proposed. Indeed, these stimulus representations may also include those corresponding to the behavioral responses they have to be associated with in the experiment, in accordance with the scalp distribution of the strongest correlations.

On the other hand, in both tasks, smaller N400s were observed for placebos than for no-pills. This may indicate that fewer of the representations activated during preconscious processing of the stimuli were inhibited by N400 processes (for support of the idea that the N400 indexes inhibitory processes, see Debruille, 2007; Debruille et al., 2008; Shang & Debruille, 2013; and Sinha et al. 2023). This could make sense, as representations of the self can only be less accurate, blurred by the belief of being under the influence of a drug that has never been experienced before. Fewer representations can then clearly mismatch this imprecise current representation of the self, in which anything can be expected. There will thus be a larger number of representations entering the content of working memory. If confirmed by further works, this would conversely raise the possibility that precise current self-representations involved in P2 processes may determine what the content of consciousness indexed by the LPP will be.

Lastly, the larger LPPs found in the cognitive test than in the social role task have to be discussed. One likely possibility is the certainty with which one can make the decision in the former than in the latter. Indeed, certainty has been found to go with larger LPPs (Hagen et al., 2006; Johnen & Harrison, 2020; Zakrzewski et al., 2023).

One limitation of the present study is that the results might be partly due to a recruitment bias in the placebo group. Indeed, a particular advertisement strategy had to be used for these participants. The advertisement stated explicitly that they would take an antipsychotic. This might have resulted in the attraction of sensation seekers. However, our results were quite different from previous ERP studies contrasting sensation seekers from non-sensation seekers (Lawson et al., 2012; Zheng et al., 2015). Furthermore, SPQ scores did not differ between groups, whereas sensation seeking usually positively correlates with schizotypy (Del Giudice et al., 2014). As the sex ratio was balanced across groups and did not differ significantly, sex-related confounds are unlikely to account for the observed effects.

To further test whether placebo effects seen in this study reflect the binding of stimulus representations with a broader set of self-representations, future research would have to find experimental conditions that manipulate these representations in a different way, such as by, for instance, having participants with a friend or with a stranger (Sinha et al., 2025; Sinha et al., 2023).

## Ethical Approval

This study was approved by the Douglas Ethics Review Board (project number: IUSMD-06-42).

## Consent to participate

Informed consent was obtained from all participants included in the study.

## Consent to publish

Participants gave explicit consent for their anonymized data to be published in the research.

## Data Availability Statement

All datasets (behavioral and EEG) can be obtained from the corresponding author upon request.

## Authors Contributions

MD reprocessed all the data collected, performed new statistical analyses and wrote the current manuscript. ID supervised part of the testing and wrote a Master’s thesis on part of the results presented here. JQ, JZ and AS contributed to the statistical analysis, the interpretation of the results, the writing of the manuscript and its corrections. SPLV contributed to the revision of the manuscript and the interpretation of the findings. JBD wrote the research project, got the funding, designed the experiment, supervised MD, ID, JQ, JZ and AS, and corrected earlier versions of this manuscript. All authors contributed to and have approved the final manuscript. We thank Ola Mohammed Ali, Gifty Asare, Timothy Hadjis, and Gabriella Vélez Largo for testing participants.

