## Supplementary materials for "Cognitive effects of placebo antipsychotics: Investigating their mechanisms using event-related potentials (ERPs) in a cognitive test and a self-referential task"

**Table of contents**

1. Text S1. Verbatim of the consent form paragraph presented to the no-pill group
2. Text S2. Verbatim of the consent form paragraph presented to the placebo group
3. Fatigue and anxiety questionnaires

3.1 Text S3. Participant's instructions for the first fatigue questionnaire

3.2 Table S1. Questions selected from the first fatigue questionnaire and used for standardization

3.3 Text S4. Participant's instructions for the second fatigue questionnaire

3.4 Table S2. Questions selected from the second fatigue questionnaire and used for standardization

3.5 Text S5. Participant's instructions for the anxiety questionnaire

3.6 Table S3. Questions selected from the anxiety questionnaire and used for standardization

4. Task instructions

4.1 Text S6. The cognitive test (=Semantic categorization task)

4.2 Text S7. The social role self-referential task

5. Stimulus timing

5.1 Figure S1. Timing of the stimulus presentations of the cognitive test

5.2 Figure S2. Timing of the stimulus presentations of the social role task

6. Text S8. Data processing
7. Figure S3. Grand average ERPs of the placebo- and no-pill-group in the cognitive test and social role task
8. Table S4. FDR-significant results of the post-hoc pairwise comparisons of the no-pill vs. placebo group run on mean voltages of the N1, P2, N400 and LPP ERPs at each electrode
9. Table S5. FDR-significant results of the post-hoc pairwise comparisons of the cognitive test vs. the social role task run on mean voltages of the LPPs at each electrode
10. Survey exploring the expectations of the effects of the antipsychotic

10.1 Text S9. Verbatim of the follow-up survey advertisement used for screening

10.2 Text S10. Verbatim of the consent form paragraph presented as part of the expectation survey

10.3 Contents of the Expectations Assessment Scale (EAS) used in the survey

10.3.1 Table S6. Effect presence subscale

10.3.2 Table S7. Effect valence subscale

10.3.3 Table S8. Effect direction subscale

10.3.4 Table S9. Effect strength subscale

10.4 Table S10. Scores obtained at the Expectations Assessment Scale (EAS)

11. Table S11. Average numbers of accepted trials and mean amplitude of the N1, P2, N400 and LPP ERPs at Cz and mean amplitude of these ERPs averaged across electrodes in each condition
12. Table S12. List of the stimuli of the cognitive test
13. Table S13. List of the stimuli of the social role task

### Text S1. Verbatim of the consent form paragraph presented to the no-pill group:

It is known that antipsychotic medications function to improve a number of psychotic symptoms. The exact mechanisms of this function remain incompletely understood. One possibility is that these medications facilitate, directly or indirectly, the mechanisms that allow us to understand unexpected information. By doing so, these medications could help patients with inaccurate beliefs (eg. delusions) to change their minds. In addition, telling participants they will take an antipsychotic when in fact they receive a placebo pill, may have an effect similar to that of these medications. Interestingly, a component of the electrical activity of the brain, termed the N400, reflects the efforts automatically deployed by the brain to understand unexpected information. As predicted, this component is smaller in patients with severe inaccurate beliefs, suggesting that their brains deploy less effort to process unexpected information. A recent study has demonstrated that even in healthy individuals, a smaller N400 accompanies the tendency towards delusional beliefs. The main goal of this study is to examine the manner in which a low dose of an antipsychotic medication, risperidone or olanzapine, as compared to placebo, influences this N400 measure and thus affects this tendency.

Testing the effect of the placebo may also provide a greater understanding of how this antipsychotic works. We thus now need a group of participants who will neither have medication nor placebo (i.e., a no pill group) and use it as a reference group. This is why you are now offered to participate in this research.

Your participation in this study would involve one session of tests lasting 3 to 4 hours and taking place at the Douglas Hospital. Your participation first consists of questionnaires that you will be requested to fill out. They include questions that aim to acquire general information such as your age, education, medical problems as well as questions related to personality characteristics, beliefs, and psychiatric symptoms. After filling out the questionnaires, you will be seated at a computer terminal. A cap that includes small metal disks that capture the electrical activity of your brain will be placed on your head. You will then have to decide if the words appearing on the screen are names of animals or not. Then, names of social roles (ex: ‘’parent’’) will appear and you will have to decide whether or not you would consider playing that role in your life. After a lunch break, you will have to make these decisions again (but this time, the medication will have had time to have its effect). The part of the experiment where you wear the cap is about 2 hours long. A follow-up debriefing session will take place at the end of the experiment.

The procedure is not painful but may be slightly uncomfortable. The only risk involved in recording the electrical activity is the rare occurrence (1% of cases) of a local allergic reaction (rash) to the adhesive on the electrodes – this carries no danger to your health. If you experience such a reaction, the recording will be stopped.

### Text S2. Verbatim of the consent form paragraph presented to the placebo group:

It is known that antipsychotic medications function to improve a number of psychotic symptoms. The exact mechanisms of this function remain incompletely understood. One possibility is that these medications facilitate, directly or indirectly, the mechanisms that allow us to understand unexpected information. By doing so, these medications could help patients with inaccurate beliefs (eg. delusions) to change their minds. Interestingly, a component of the electrical activity of the brain, termed the N400, reflects the efforts automatically deployed by the brain to understand unexpected information. As predicted, this component is smaller in patients with severe inaccurate beliefs, suggesting that their brains deploy less effort to process unexpected information. A recent study has demonstrated that even in healthy individuals, a smaller N400 accompanies the tendency towards delusional beliefs. The main goal of this study is to examine the manner in which a low dose of an antipsychotic medication, risperidone or olanzapine, influences this N400 measure and thus affects this tendency. Moreover, it includes a number of other tests that will be used to make sure that we are actually focusing on the tendency towards delusional belief rather than on another tendency.

Your participation in this study would involve one session of tests lasting 3 to 4 hours and taking place at the Douglas Hospital. It consists first of questionnaires that you will be requested to fill out. They include questions that aim to acquire general information such as your age, education, medical problems as well as questions related to personality characteristics, beliefs, and psychiatric symptoms. After filling the questionnaires having you will be sited at a computer terminal. A cap that includes small metal disks that capture the electrical activity of your brain will be placed on your head. Once this will be done, you will be given 2.5 milligrams of an active medication, risperidone or olanzapine. You will then have to decide if the words appearing on the screen are names of animals or not. Then, names of social roles (ex: ‘’parent’’) will appear and you will have to decide whether or not you would consider playing that role in your life. After a lunch break, you will have to make these decisions again (but this time, the medication will have had time to have its effect). The part of the experiment where you wear the cap is about 2 hours long.

The procedure is not painful but may be slightly uncomfortable. The only risk involved in recording the electrical activity is the rare occurrence (1% of cases) of a local allergic reaction (rash) to the adhesive on the electrodes – this carries no danger to your health. If you experience such a reaction, the recording will be stopped. Risperidone or olanzapine is a drug that is widely used in clinical practice by thousands of patients and is approved by the Food and Drug Administration. The 2.5 milligram dose you will be taking is almost the lowest dose that is given to adults. There are several possible side-effects associated with this medication, although it is extremely unlikely that they could occur at the dose involved in this study. Also, note that all studies that examined side-effects looked at repeated administrations of the drug, rather than at single dose. The adverse effects of this dose of risperidone or olanzapine that you might experience are somnolence, dry mouth, light-headedness, constipation, increased appetite, stomach upset (nausea/indigestion), restlessness, sense of muscle weakness (with no actual loss of strength), insomnia, and muscle stiffness. However, mild drowsiness is the only adverse effect that is likely to occur.

### Fatigue and anxiety questionnaires

Over the entire course of the study, two fatigue and anxiety questionnaires were used in succession to measure the energy and anxiety levels of the participants. In order to unify the analysis, we selected, from both questionnaires, the questions that reflected the current energy and anxiety levels of the participants and standardized them using the percentage maximum possible (POMP) method (Cohen et al., 1999). Each POMP score of each participant equal 100 * (raw - min) / (max - min), with raw = original mean score computed with the score for each of the selected questions with valid values, min = minimum possible value, and max = maximum possible value. For example, we selected 14 questions for the STAI questionnaire, all of which used a 4-point Likert scale. A participant had a raw score on the STAI questionnaire of 50. So, POMP for this participant = 100 * (50 – 14 x 1) / (14 x 4 – 14 x 1).

#### Text S3. Participant's instructions for the first fatigue questionnaire:

Please answer each question on a scale of 1 to 10 to indicate how you currently feel - that is, in the present moment. Answer ALL items even if unsure of your answer. When you have finished, check over each one to make sure you have answered them all (1-Lowest; 10-Extremely high).

#### Table S1. Questions selected from the first fatigue questionnaire and used for standardization.

|  | **1** | **2** | **3** | **4** | **5** | **6** | **7** | **8** | **9** | **10** |
| --- | --- | --- | --- | --- | --- | --- | --- | --- | --- | --- |
| How is your level of energy? | **○** | **○** | **○** | **○** | **○** | **○** | **○** | **○** | **○** | **○** |
| How sleepy are you? | **○** | **○** | **○** | **○** | **○** | **○** | **○** | **○** | **○** | **○** |

#### Text S4. Participant's instructions for the second fatigue questionnaire:

The following ten statements refer to how you currently feel. Please select the answer to each question that is applicable to you. Please give an answer to each question (1-Never; 5-Always).

#### Table S2. Questions selected from the second fatigue questionnaire and used for standardization.

|  | **1** | **2** | **3** | **4** | **5** |
| --- | --- | --- | --- | --- | --- |
| I am bothered by fatigue; | **○** | **○** | **○** | **○** | **○** |
| Physically, I feel exhausted | **○** | **○** | **○** | **○** | **○** |
| I feel no desire to do anything | **○** | **○** | **○** | **○** | **○** |
| Mentally, I feel exhausted | **○** | **○** | **○** | **○** | **○** |

#### Text S5. Participant's instructions for the anxiety questionnaire:

A number of statements which people have used to describe themselves are given below. Read each statement and then select the appropriate number next to the statement to indicate how you feel right now, that is, at this moment. There are no right or wrong answers. Do not spend too much time on any one statement but give the answer which seems to describe your present feelings best (1-No at all; 4-Very much so).

#### Table S3. Questions selected from the anxiety questionnaire and used for standardization.

|  | **1** | **2** | **3** | **4** |
| --- | --- | --- | --- | --- |
| I feel calm | **○** | **○** | **○** | **○** |
| I feel secure | **○** | **○** | **○** | **○** |
| I am tense | **○** | **○** | **○** | **○** |
| I feel at ease | **○** | **○** | **○** | **○** |
| I feel upset | **○** | **○** | **○** | **○** |
| I am presently worrying over possible misfortunes | **○** | **○** | **○** | **○** |
| I feel comfortable | **○** | **○** | **○** | **○** |
| I feel self-confident | **○** | **○** | **○** | **○** |
| I feel nervous | **○** | **○** | **○** | **○** |
| I am jittery | **○** | **○** | **○** | **○** |
| I am relaxed | **○** | **○** | **○** | **○** |
| I feel content | **○** | **○** | **○** | **○** |
| I am worried | **○** | **○** | **○** | **○** |
| I feel pleasant | **○** | **○** | **○** | **○** |

Cohen, P., Cohen, J., Aiken, L. S., & West, S. G. (1999). The problem of units and the circumstance for POMP. *Multivariate behavioral research,* 34(3), 315-346. <https://doi.org/10.1207/s15327906mbr3403_2>

### Task instructions

The experiment will last approximately 20 minutes. You are free to refuse or stop participating at any time without being asked for a reason and without any negative consequences.

To avoid eye movements during the experiment, focus only on the center of the screen. The “+” indicates that center. Your gaze should remain there at all times unless you are instructed to blink.

It is important to stay relaxed and still throughout the experiment. Please avoid head-, eye-, facial-, or body-movement.

This pertains most importantly to blinking. Please blink ONLY when you are asked to. This will occur approximately every 30 seconds. Use this time, and only this time, to blink, swallow your saliva, or move any body parts as needed.

#### Text S6. The cognitive test (=Semantic categorization task):

In this task, you will see the serial presentation of two words.

In most trials, the first of these words the prime ‘ANIMAL’. It is followed by a ‘TARGET’ word that matches the prime category (eg. an animal name, eg., dog) or does NOT match the prime category (eg., an object name, eg., couch).

Respond as accurately and as fast as possible by pressing the 1 button on the keyboard if the two words match, and the 2 button if the words do not match.

However, in some trials, the first of these words is the prime ‘INACTION’ and is followed by a matching or mismatching target word. In this case, DO NOT respond on the keyboard.

The experiment will automatically proceed to the next trial.

#### Text S7. The social role self-referential task:

In this task, you will see names of social roles (which could be any pattern of behavior, for example a parent, superman or drug addict).

For each name, press the YES button on the response pad if you COULD CONSIDER YOURSELF PLAYING THIS ROLE AT ANY MOMENT IN YOUR LIFE. If not, press the NO button.

Consider the role in your respective gender, despite the gender that may appear on screen. You may respond even after the names have disappeared.

Each name will be presented for ½ a second only and a cross will appear on the screen after your response. Please concentrate on this cross. The experiment will proceed to the next name automatically.

### Stimulus timing

#### Figure S1. Timing of the stimulus presentations of the cognitive test.

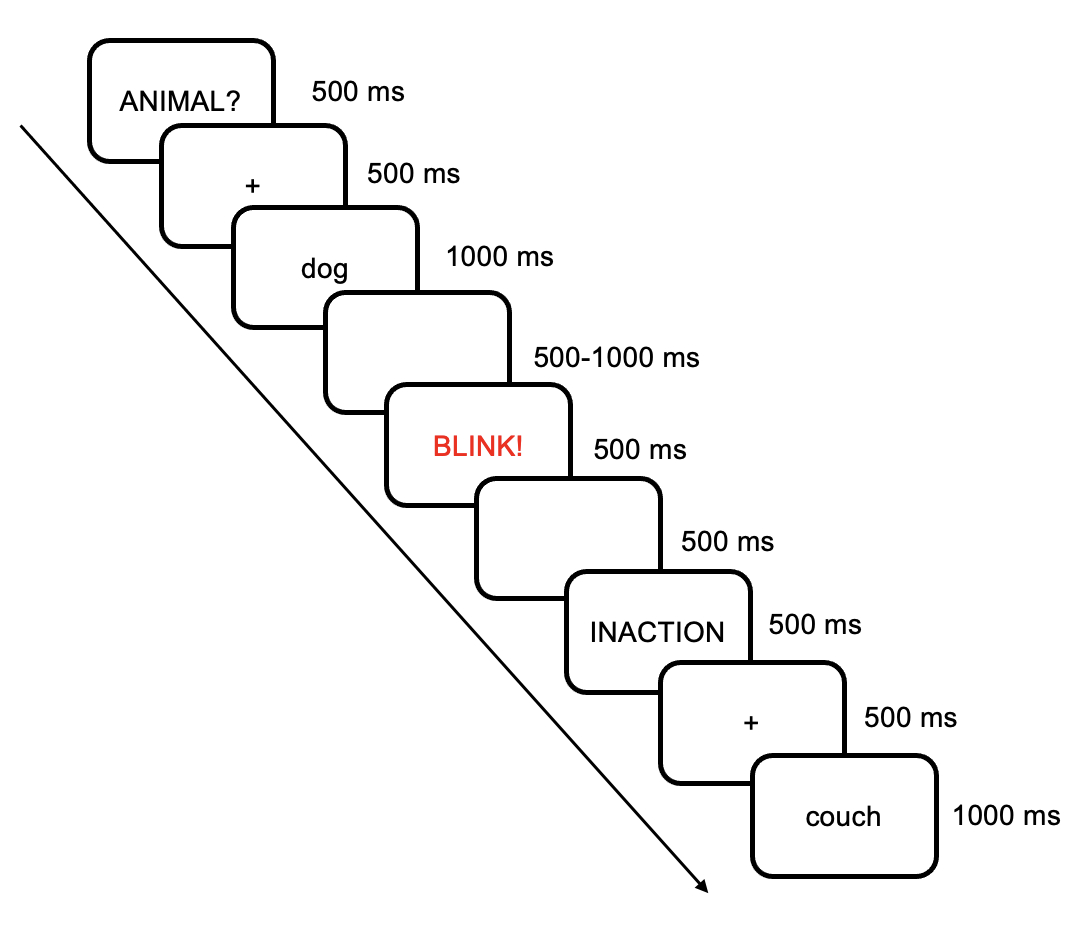

#### Figure S2. Timing of the stimulus presentations of the social role task.

All participants were presented with the same social role stimuli. However, there were differences in the duration of the stimuli, as shown in the figure below. For timing 1, each role was presented for 500 ms, followed immediately by a fixation cross, which lasted for a randomized duration comprised between 1500 to 2500 ms. For every five roles, the fixation cross was replaced by a ‘BLINK!’ stimulus, which lasted for 500 ms. For timing 2, each role was presented for 1800 ms, immediately followed by a ‘BLINK!’ stimulus, which lasted for 1000 ms, and was then replaced by a fixation cross, which lasted for a randomized duration comprised between 300 to 1000 ms. For timing 3, each role was presented for 1800 ms, immediately followed by a ‘BLINK!’ stimulus, which lasted for 500 ms and was then replaced by a fixation cross, which lasted for a randomized duration comprised between 800 to 1500 ms. All 43 participants in the no-pill group used timing 3. Of the placebo group, 27 participants used timing 1, 9 participants used timing 2, and 4 participants used timing 3. No significant differences were found in the P2 and LPP ERPs between the no-pill and placebo groups in this self-referential social-role task. On the other hand, the group differences found on the amplitudes of N400s cannot be due to these timing differences as, even with the fastest timing, there was still 2 seconds for participants to make their decision while the mean RT was 1017 ms (SD = 164).

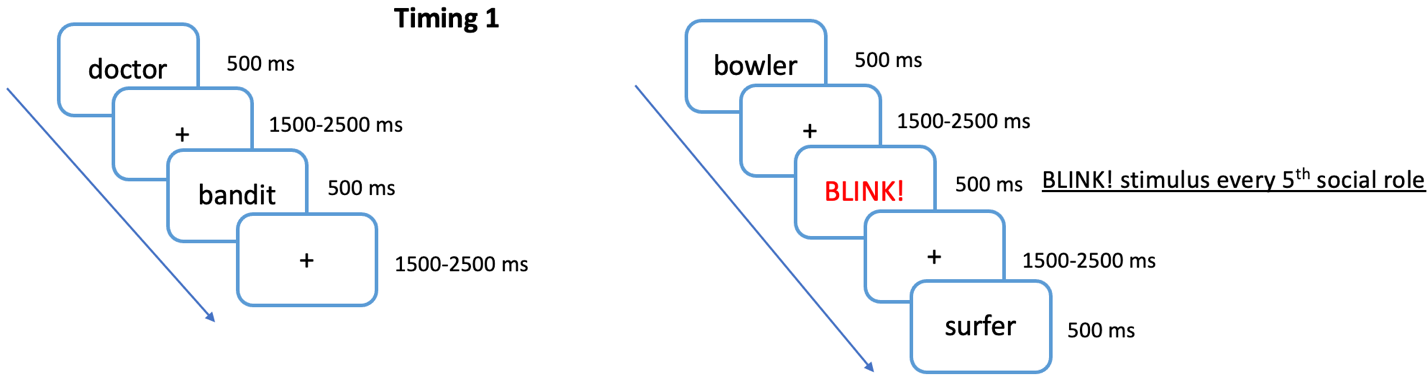

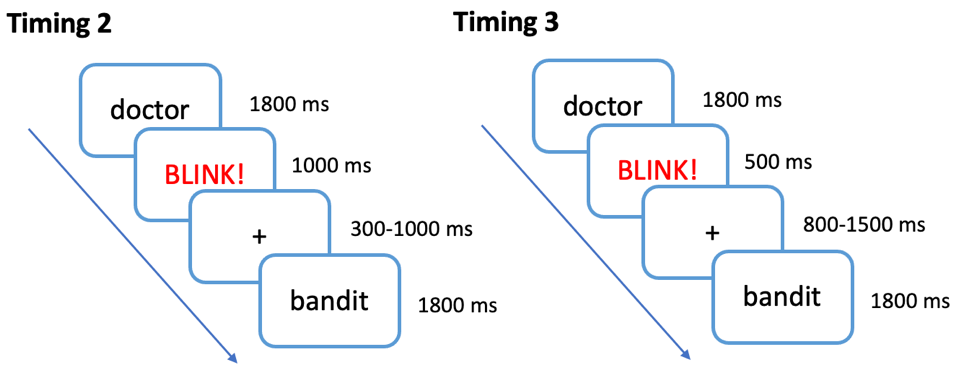

### Text S8. Data processing

An independent component analysis (ICA) was used to remove artifacts, including but not limited to blinks, eye movements and myograms (Bell and Sejnowski, 1995). The infomax algorithm ICA was performed on a copy of the continuous EEG that was high-pass filtered at 1 Hz and low-pass filtered at 30 Hz. For that, the resulting ICA weight matrix and sphering matrix were then applied to the continuous 0.1-30 Hz filtered EEG by using the ICLabel EEGLAB extension to signal artifactual independent components (ICs) that had more than 20% of chance of being muscle activity or more than 8% of chance of being eye movements (Cline et al., 2021). These ICs were then systematically subtracted from this continuous 0.1-30 Hz filtered EEG, as in Finke et al. (2016), Goregliad et al. (2016) and Markey et al. (2019). For both cognitive and social role tasks, only trials including behavioral responses performed between 300 ms and 2500 ms post-onset were retained to eliminate trials to which participants did not pay enough attention to, were too hesitant, or provided rash responses. The EEG epochs for those retained trials were taken from 200 ms pre-stimulus to 1000 ms post-stimulus. Their baselines were set by computing the mean voltage in the -200 to 0 ms time window for each electrode and by subtracting this mean value from each point of the -200 to 1000 ms epoch. Trials with voltages whose amplitude exceeded -/+100 μV at one or more of the four frontal electrodes (Fp1/2, F7/8) or exceeded -/+75 μV at one or more of the other 24 electrode sites were rejected. Trials with one or more flat lines lasting longer than 100 ms were also excluded.

Bell AJ, Sejnowski TJ. An information-maximization approach to blind separation and blind deconvolution. Neural Comput. 1995;7(6):1129-59. doi: 10.1162/neco.1995.7.6.1129. PubMed PMID: 7584893.

Cline CC, Lucas MV, Sun Y, Menezes M, Etkin A, editors. Advanced artifact removal for automated TMS-EEG data processing. 2021 10th International IEEE/EMBS Conference on Neural Engineering (NER); 2021: IEEE.

Finke M, Büchner A, Ruigendijk E, Meyer M, Sandmann P. On the relationship between auditory cognition and speech intelligibility in cochlear implant users: An ERP study. Neuropsychologia. 2016;87:169-81. Epub 20160519. doi: 10.1016/j.neuropsychologia.2016.05.019. PubMed PMID: 27212057.

Goregliad Fjaellingsdal T, Ruigendijk E, Scherbaum S, Bleichner MG. The N400 Effect during Speaker-Switch-Towards a Conversational Approach of Measuring Neural Correlates of Language. Front Psychol. 2016;7:1854. Epub 20161128. doi: 10.3389/fpsyg.2016.01854. PubMed PMID: 27965604; PubMed Central PMCID: PMC5124707.

Markey PS, Jakesch M, Leder H. Art looks different - Semantic and syntactic processing of paintings and associated neurophysiological brain responses. Brain Cogn. 2019;134:58-66. Epub 20190528. doi: 10.1016/j.bandc.2019.05.008. PubMed PMID: 31151085.

### Figure S3. Grand average ERPs of the placebo- and no-pill-group in the cognitive test and social role task

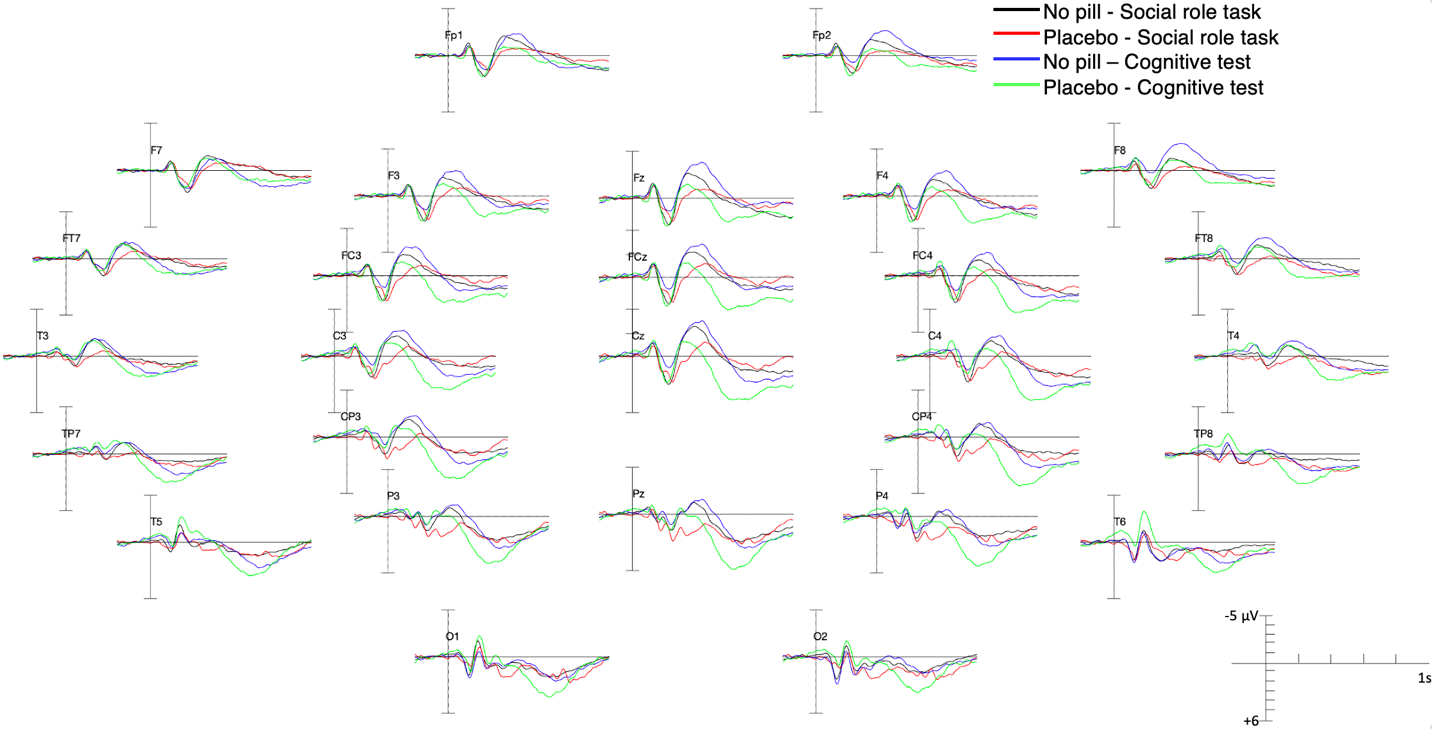

*Grand average ERPs. Black lines correspond to the no-pill participants (N =43) in the social role task, red lines correspond to the placebo participants (N = 40) in this task, blue lines correspond to the no-pill participants in the cognitive test, green lines correspond to the placebo participants in that test.*

### Table S4. FDR-significant results of the post-hoc pairwise comparisons of the no-pill vs. placebo group run on mean voltages of the N1, P2, N400 and LPP ERPs at each electrode

| N1 in COG | | P2 in COG | | N400 in COG | | LPP in MM | | LPP in M | |
| --- | --- | --- | --- | --- | --- | --- | --- | --- | --- |
| T6 | 4.1x10^-6^ | FP2 | 0.001 | Fp2 | 5.8x10^-4^ | Fp2 | 0.008 | Fp2 | 0.039 |
| T5 | 0.007 | F8 | 0.001 | F8 | 4x10^-4^ | F8 | 1x10^-4^ | F8 | 0.007 |
| O2 | 0.015 | FZ | 0.012 | Fz | 0.003 | Fz | 0.005 | Cz | 0.025 |
| O1 | 0.044 | CZ | 0.03 | Cz | 0.011 | Cz | 8.1x10^-4^ | Pz | 0.014 |
|  |  | F4 | 0.007 | Pz | 0.034 | Pz | 4.4x10^-4^ | P4 | 0.016 |
|  |  | FC3 | 0.02 | F4 | 0.009 | P4 | 2.4x10^-4^ | Ft8 | 0.034 |
|  |  | FCZ | 0.006 | F3 | 0.029 | P3 | 0.003 | Fc4 | 0.036 |
|  |  | TP8 | 0.002 | Ft8 | 0.027 | T6 | 0.036 | Fcz | 0.02 |
|  |  |  |  | Fc4 | 0.021 | T5 | 0.059 | C4 | 0.037 |
|  |  |  |  | Fc3 | 0.013 | T4 | 1.8x10^-4^ | C3 | 0.046 |
|  |  |  |  | Fcz | 0.006 | F4 | 0.001 | Tp8 | 0.036 |
|  | |  |  |  |  | F3 | 0.026 | Cp4 | 0.013 |
|  | |  |  |  |  | Ft8 | 4.5x10^-4^ | Cp3 | 0.034 |
|  | |  |  |  |  | Fc4 | 6.2x10^-4^ | O2 | 0.031 |
|  | |  |  |  |  | Fc3 | 0.003 |  |  |
|  | |  |  |  |  | Fcz | 8.8x10^-4^ |  |  |
|  | |  |  |  |  | C4 | 7.5x10^-4^ |  |  |
|  | |  |  |  |  | C3 | 0.002 |  |  |
|  | |  |  |  |  | Tp8 | 3.9x10^-4^ |  |  |
|  | |  |  |  |  | Tp7 | 0.024 |  |  |
|  | |  |  |  |  | Cp4 | 2.2x10^-4^ |  |  |
|  | |  |  |  |  | Cp3 | 0.002 |  |  |
|  | |  |  |  |  | O2 | 7.1x10^-4^ |  |  |
|  | |  |  |  |  | O1 | 0.013 |  |  |

*COG: cognitive test; MM: mismatch condition of the cognitive test; M: match condition of the cognitive test.*

### Table S5. FDR-significant results of the post-hoc pairwise comparisons of the cognitive test vs. the social role task run on mean voltages of the LPPs at each electrode

| LPP in MM of two tasks | | LPP in M of two tasks | |
| --- | --- | --- | --- |
| Fp1 | 0.057 | F7 | 0.001 |
| F7 | 0.003 | Cz | 2.8x10^-4^ |
| Cz | 4.5x10^-4^ | Pz | 0.001 |
| Pz | 0.025 | P4 | 3.9x10^-5^ |
| P4 | 0.004 | P3 | 0.001 |
| P3 | 0.022 | T6 | 4.8x10^-6^ |
| T6 | 0.001 | T5 | 8x10^-6^ |
| T5 | 7.3x10^-4^ | T4 | 1.2x10^-4^ |
| T4 | 0.039 | T3 | 7.2x10^-4^ |
| T3 | 0.002 | F4 | 0.053 |
| F4 | 0.014 | Ft8 | 0.011 |
| F3 | 0.019 | Ft7 | 0.002 |
| Ft7 | 0.001 | Fc4 | 4.3x10^-4^ |
| Fc4 | 3.9x10^-4^ | Fc3 | 0.004 |
| Fc3 | 0.004 | Fcz | 0.027 |
| Fcz | 0.005 | C4 | 7.9x10^-6^ |
| C4 | 7.1x10^-4^ | C3 | 7.6x10^-5^ |
| C3 | 0.001 | Tp8 | 7.1x10^-6^ |
| Tp8 | 0.002 | Tp7 | 7.1x10^-6^ |
| Tp7 | 1.3x10^-4^ | Cp4 | 1x10^-6^ |
| Cp4 | 3.2x10^-4^ | Cp3 | 1.3x10^-4^ |
| Cp3 | 0.004 | O2 | 0.056 |
| O1 | 0.041 | O1 | 0.011 |

*MM: mismatch condition; M: match condition.*

### Survey exploring the expectations of the effects of the antipsychotic

#### Text S9. Verbatim of the follow-up survey advertisement used for screening:

“But first, we have to ask you whether you would participate in a follow-up study that will be evaluating the effect of antipsychotic medications. As such, a low dose of an antipsychotic (risperidone or olanzapine) will be given to you. In addition, an electrode cap will be placed on your head to record the brain’s electrical activity (EEG) while you perform simple tasks on a computer. This follow-up brain study is separate from the current survey so keep in mind that you might not be offered to participate in it. This study would take place at the Douglas Mental Health Hospital in Verdun and would last approximately 3-4 hours. You would be compensated $13 per hour and you could expect to receive at least $45 (+ a full reimbursement of travel expenses).”

#### Text S10. Verbatim of the consent form paragraph presented as part of the expectation survey:

*Assume you are participating in a study where you are given a single dose of antipsychotic medication olanzapine. Consider the following information:*

It is known that antipsychotic medications function to improve a number of psychotic symptoms. The exact mechanisms of the function remain incompletely understood. One possibility is that these medications facilitate, directly or indirectly, the mechanisms that allow us to understand unexpected information. By doing so, these medications could help patients with inaccurate beliefs (e.g., delusions) to change their minds.

Olanzapine is a drug that is widely used in clinical practice by thousands of patients and is approved by the Food and Drug Administration. The 2.5 milligram dose you will be taking is almost the lowest dose that is given to adults. There are several possible side effects associated with this medication, although it is extremely unlikely that they could occur at the dose involved in this study. Also, note that all studies that examined side effects looked at repeated administration of the drug, rather than at a single dose.

The adverse effects that you might experience are sleepiness and drowsiness, dry mouth, light-headedness, constipation, increased appetite, stomach upset (nausea / indigestion), restlessness, sense of muscle weakness (with no actual loss of strength), insomnia, and muscle stiffness. However, mild drowsiness is the only adverse effect that is likely to occur.

#### Contents of the Expectations Assessment Scale (EAS) used in the survey

##### **Table S6. Effect presence subscale.**

Please use the following scale to indicate whether you think an antipsychotic would affect certain aspects of your thinking and personality to a large extent or not at all. When answering this question, please think about whether an antipsychotic would have an effect or not.

0 – Not at all;

1 – To a small extent;

2 – To some extent;

3 – To a moderate extent;

4 – To a great extent;

5 – To a very great extent;

|  | **0** | **1** | **2** | **3** | **4** | **5** |
| --- | --- | --- | --- | --- | --- | --- |
| Cognitive functions (attention, memory, visual perception, information processing and reasoning) | **○** | **○** | **○** | **○** | **○** | **○** |
| Memory | **○** | **○** | **○** | **○** | **○** | **○** |
| Concentration | **○** | **○** | **○** | **○** | **○** | **○** |
| Distractibility | **○** | **○** | **○** | **○** | **○** | **○** |
| Reasoning ability | **○** | **○** | **○** | **○** | **○** | **○** |
| Multitasking ability | **○** | **○** | **○** | **○** | **○** | **○** |
| Performance in everyday tasks (e.g., driving, remembering important days, managing finances, etc.) | **○** | **○** | **○** | **○** | **○** | **○** |
| Thought process | **○** | **○** | **○** | **○** | **○** | **○** |
| Levels of energy | **○** | **○** | **○** | **○** | **○** | **○** |
| Anxiety | **○** | **○** | **○** | **○** | **○** | **○** |
| Impulsivity | **○** | **○** | **○** | **○** | **○** | **○** |
| Spontaneity | **○** | **○** | **○** | **○** | **○** | **○** |
| Creativity | **○** | **○** | **○** | **○** | **○** | **○** |
| Perception of pain | **○** | **○** | **○** | **○** | **○** | **○** |
| Feeling of nausea | **○** | **○** | **○** | **○** | **○** | **○** |
| Sleepiness | **○** | **○** | **○** | **○** | **○** | **○** |
| Movement and motion | **○** | **○** | **○** | **○** | **○** | **○** |
| Appetite | **○** | **○** | **○** | **○** | **○** | **○** |
| Sexual function | **○** | **○** | **○** | **○** | **○** | **○** |

*Note*. Items adapted from Rabipour et al. (2018): Cognitive functions, Memory, Concentration, Distractibility, Reasoning ability, Multitasking ability, Performance in everyday tasks.

##### Table S7. Effect valence subscale.

Please use the following scale to indicate whether you think an antipsychotic would affect certain aspects of your thinking and personality very positively or very negatively. When answering this question, please think about what kind of effect an antipsychotic would have.

1 – Very negatively (score -3);

2 – Fairly negatively (score -2);

3 – Somewhat negatively (score -1);

4 – I have absolutely no expectations (score 0);

5 – Somewhat positively (score 1);

6 – Fairly positively (score 2);

7 – Very positively (score 3);

|  | **1** | **2** | **3** | **4** | **5** | **6** | **7** |
| --- | --- | --- | --- | --- | --- | --- | --- |
| Cognitive functions (attention, memory, visual perception, information processing and reasoning) | **○** | **○** | **○** | **○** | **○** | **○** | **○** |
| Memory | **○** | **○** | **○** | **○** | **○** | **○** | **○** |
| Concentration | **○** | **○** | **○** | **○** | **○** | **○** | **○** |
| Distractibility | **○** | **○** | **○** | **○** | **○** | **○** | **○** |
| Reasoning ability | **○** | **○** | **○** | **○** | **○** | **○** | **○** |
| Multitasking ability | **○** | **○** | **○** | **○** | **○** | **○** | **○** |
| Performance in everyday tasks (e.g., driving, remembering important days, managing finances, etc.) | **○** | **○** | **○** | **○** | **○** | **○** | **○** |
| Thought process | **○** | **○** | **○** | **○** | **○** | **○** | **○** |
| Appetite | **○** | **○** | **○** | **○** | **○** | **○** | **○** |
| Sexual function | **○** | **○** | **○** | **○** | **○** | **○** | **○** |

*Note*. Items adapted from Rabipour et al. (2018): Cognitive functions, Memory, Concentration, Distractibility, Reasoning ability, Multitasking ability, Performance in everyday tasks.

##### Table S8. Effect direction subscale.

A single minimal dose of an antipsychotic would make me

1 – Significantly less (score -3);

2 – Much less (score -2);

3 – Somewhat less (score -1);

4 – I have absolutely no expectations (score 0);

5 – Somewhat more (score 1);

6 – Much more (score 2);

7 – Significantly more (score 3);

|  | **1** | **2** | **3** | **4** | **5** | **6** | **7** |
| --- | --- | --- | --- | --- | --- | --- | --- |
| Tired | **○** | **○** | **○** | **○** | **○** | **○** | **○** |
| Anxious | **○** | **○** | **○** | **○** | **○** | **○** | **○** |
| Impulsive | **○** | **○** | **○** | **○** | **○** | **○** | **○** |
| Spontaneous | **○** | **○** | **○** | **○** | **○** | **○** | **○** |
| Creative | **○** | **○** | **○** | **○** | **○** | **○** | **○** |
| Able to feel pain | **○** | **○** | **○** | **○** | **○** | **○** | **○** |
| Nauseous | **○** | **○** | **○** | **○** | **○** | **○** | **○** |
| Sleepy | **○** | **○** | **○** | **○** | **○** | **○** | **○** |
| Agitated | **○** | **○** | **○** | **○** | **○** | **○** | **○** |

##### Table S9. Effect strength subscale.

Please use the following scale to indicate to what extent you think an antipsychotic would make you feel

0 – Not at all;

1 – To a small extent;

2 – To some extent;

3 – To a moderate extent;

4 – To a great extent;

5 – To a very great extent;

|  | **0** | **1** | **2** | **3** | **4** | **5** |
| --- | --- | --- | --- | --- | --- | --- |
| Sad | **○** | **○** | **○** | **○** | **○** | **○** |
| Joyful | **○** | **○** | **○** | **○** | **○** | **○** |
| Emotionally blunted | **○** | **○** | **○** | **○** | **○** | **○** |
| Excited | **○** | **○** | **○** | **○** | **○** | **○** |
| Weird | **○** | **○** | **○** | **○** | **○** | **○** |
| Defenceless | **○** | **○** | **○** | **○** | **○** | **○** |
| Confused | **○** | **○** | **○** | **○** | **○** | **○** |
| Psychotic (having difficulties determining what is real and what is not, having false beliefs, seeing or hearing things that others do not see or hear) | **○** | **○** | **○** | **○** | **○** | **○** |

#### Table S10. Scores obtained at the Expectations Assessment Scale (EAS)

| *Within-group comparison (Wilcoxon signed-rank test) on the EAS items before and after the consent form paragraph presentation for participants (N = 52)* | | | | | |
| --- | --- | --- | --- | --- | --- |
| EAS Question type | EAS Item | Median response before reading the consent form paragraph | Median response after reading the consent form paragraph | *p*  (1-tailed) | *η^2^* |
| Effect Presence^a^ | Cognitive Functions | To a moderate extent | To some extent | 0.006 | 0.11 |
|  | Anxiety | To a moderate extent | To a small extent | 0.0000015 | 0.38 |
|  | Concentration | To a moderate extent | To some extent | 0.00015 | 0.23 |
|  | Creativity | To some extent | To a small extent | 0.00002 | 0.29 |
|  | Distractibility | To some extent | To some extent | 0.00005 | 0.25 |
|  | Energy | To a moderate extent | To some extent | 0.0005 | 0.18 |
|  | Impulsivity | To a moderate extent | To a small extent | 0.0000005 | 0.41 |
|  | Memory | To some extent | To a small extent | 0.004 | 0.12 |
|  | Multitasking ability | To some extent | To a small extent | 0.003 | 0.13 |
|  | Performance in everyday tasks | To some extent | To a small extent | 0.0285 | 0.06 |
|  | Sexual function | To some extent | To a small extent | 0.0015 | 0.16 |
|  | Spontaneity | To a moderate extent | To a small extent | 4.9 x 10^-7^ | 0.42 |
|  | Thought process | To a moderate extent | To some extent | 0.00035 | 0.20 |
|  | Perception of Pain | To a small extent | Not at all | 0.002 | 0.14 |
| Effect Valence^b^ | Memory | No expectations | No expectations | 0.036 | 0.057 |
|  | Appetite | Somewhat negatively | Somewhat negatively | 0.029 | 0.063 |
|  | Sex Function | Somewhat negatively | No expectations | 0.0004 | 0.20 |
| Effect Direction^c^ | Able to Feel Pain | No expectations | No expectations | 0.005 | 0.12 |
|  | Tired | No expectations | Somewhat more | 0.0004 | 0.20 |
|  | Agitated | Somewhat less | No expectations | 0.01 | 0.095 |
| Effect Strength^d^ | Sad | To a small extent | To a small extent | 0.022 | 0.071 |
|  | Confused | To a small extent | Not at all | 0.022 | 0.071 |
|  | Emotionally Blunted | To some extent | To a small extent | 0.00015 | 0.23 |
|  | Weird | To some extent | To a small extent | 0.001 | 0.18 |
| ^a^ Effect Presence Question type: Do you think an antipsychotic would affect your [X] to a large extent or not at all?  ^b^ Effect Valence Question type: Do you think an antipsychotic would affect your [X] very positively or very negatively?  ^c^ Effect Direction Question type: Do you think an antipsychotic would make you feel more or less [X]?  ^d^ Effect Strength Question type: Indicate to what extent you think an antipsychotic would make you feel [X] | | | | | |

### Table S11. Average numbers of accepted trials and mean amplitude of the N1, P2, N400 and LPP ERPs at Cz and mean amplitude of these ERPs averaged across electrodes in each condition.

|  |  | Placebo group | | | | | | | | | | No-pill group | | | | | |
| --- | --- | --- | --- | --- | --- | --- | --- | --- | --- | --- | --- | --- | --- | --- | --- | --- | --- |
|  |  | Social role task | | | | Cognitive test | | | | | | Social role task | | | Cognitive test | | |
|  |  | Acceptance | | Rejection | | Match | | | Mismatch | Acceptance | | | Rejection | | | Match | Mismatch |
| Numbers of trials accepted | *M* (SD) | 74.3 (31.6) | | 114.8 (31.2) | | 51.7 (6.3) | | | 54.3 (5.5) | 68.7 (28.4) | | | 110.8 (36.1) | | | 44.3 (9.8) | 47.2 (6.7) |
| Averaged N1 at 4 electrodes (µV) | *M* (SD) | | 0.2 (2.3) | | 0.3 (2.0) | | -1.1 (2.5) | -0.9 (2.4) | | | 0.05 (1.9) | | | -0.05 (1.9) | 0.3 (2.2) | | 0.5 (2.1) |
|  | Range | | -6.5 - 5.6 | | -5.2 - 5.1 | | -8.7 - 6.0 | -7.1 - 3.7 | | | -5.8 - 4.1 | | | -6.3 - 5.0 | -6.9 – 7.5 | | -8.5 – 6.1 |
| P2 at Cz (µV) | *M* (SD) | 2.0 (2.8) | | 2.0 (2.6) | | 1.8 (2.7) | | | 2.1 (2.5) | 2.1 (2.4) | | | 2.3 (2.4) | | | 0.7 (2.8) | 0.7 (2.7) |
|  | Range | -3.7 − 7.8 | | -3.9 − 8.0 | | -5.6 −8.9 | | | -3.5 − 7.8 | -3.2 − 7.3 | | | -4.7 − 8.0 | | | -6.9 − 6.7 | -7.2 − 6.4 |
| Averaged P2 at 24 electrodes (µV) | *M* (SD) | 1.4 (2.1) | | 1.4 (2.1) | | 1.0 (2.2) | | | 1.1 (2.1) | 1.4 (1.9) | | | 1.4 (1.9) | | | 0.6 (2.1) | 0.6 (2.1) |
|  | Range | -5.2 − 7.8 | | -5.4 − 9.4 | | -5.6 − 8.9 | | | -5.5 − 7.8 | -4.1 − 8.1 | | | -4.7 − 8.0 | | | -6.9 − 9.2 | -7.2 − 12.3 |
| N400 at Cz (µV) | *M* (SD) | -0.1 (3.1) | | -0.5 (2.9) | | 0.4 (4.4) | | | -1.4 (3.4) | -1.9 (3.3) | | | -2.2 (3.3) | | | -2.2 (4.0) | -3.2 (4.1) |
|  | Range | -8.5 − 7.1 | | -7.0 − 7.3 | | -11.2 − 9.2 | | | -9.8 − 5.5 | -10.0-5.1 | | | -8.7 − 6.9 | | | -9.8 − 5.2 | -10.5 − 4.5 |
| Averaged N400 at 28 electrodes (µV) | *M* (SD) | 0.5 (2.4) | | 0.3 (2.4) | | 0.4 (3.0) | | | -0.8 (2.5) | -0.5 (2.4) | | | -0.8 (2.5) | | | -0.8 (2.9) | -1.6 (2.9) |
|  | Range | -8.5 − 8.7 | | -8.2 − 8.6 | | -11.2 − 10.8 | | | -9.8 − 7.5 | -10.3 − 7.8 | | | -10.1 − 7.2 | | | -9.8 − 10.5 | -10.5 − 6.6 |
| LPP at Cz (µV) | *M* (SD) | 0.7 (3.6) | | 0.4 (3.9) | | 4.0 (4.2) | | | 4.2 (4.2) | 1.0 (4.8) | | | 0.8 (4.9) | | | 1.6 (5.1) | 0.8 (4.7) |
|  | Range | -6.6 − 8.4 | | -6.8 − 10.4 | | -4.6 − 14.7 | | | -4.9 − 11.6 | -12.8 − 11.5 | | | -12.8 − 9.0 | | | -12.2 − 11.4 | -7.9 − 10.6 |
| Averaged LPP at 28 electrodes (µV) | *M* (SD) | 0.9 (2.7) | | 0.8 (2.8) | | 2.6 (3.1) | | | 2.7 (3.2) | 0.9 (3.4) | | | 0.7 (3.3) | | | 1.4 (3.4) | 0.8 (3.3) |
|  | Range | -7.5 − 9.5 | | -9.1 − 12.2 | | -7.0 − 17.3 | | | -7.4 − 14.1 | -12.8 − 14.1 | | | -12.8 − 13.6 | | | -12.2 − 12.7 | -11.5 − 15.3 |

### Table S12. List of the stimuli of the cognitive test.

| INACTION | closet |
| --- | --- |
| ANIMAL? | basin |
| INACTION | shirt |
| ANIMAL? | alligator |
| ANIMAL? | puma |
| ANIMAL? | panther |
| ANIMAL? | cloth |
| INACTION | parachute |
| ANIMAL? | octopus |
| INACTION | fleece |
| ANIMAL? | alarm |
| INACTION | bus |
| ANIMAL? | rabbit |
| INACTION | leopard |
| ANIMAL? | swallow |
| INACTION | deer |
| ANIMAL? | whale |
| ANIMAL? | lace |
| ANIMAL? | viper |
| ANIMAL? | lizard |
| ANIMAL? | pigeon |
| ANIMAL? | wasp |
| ANIMAL? | keyboard |
| ANIMAL? | helmet |
| INACTION | cylinder |
| INACTION | bicycle |
| ANIMAL? | spoon |
| ANIMAL? | bill |
| INACTION | ski |
| ANIMAL? | camel |
| ANIMAL? | earthworm |
| ANIMAL? | satchel |
| ANIMAL? | match |
| ANIMAL? | dummy |
| INACTION | flea |
| ANIMAL? | boa |
| INACTION | hedge |
| ANIMAL? | cod |
| ANIMAL? | boot |
| ANIMAL? | parrot |
| INACTION | icicle |
| INACTION | jelly-fish |
| ANIMAL? | eagle |
| ANIMAL? | ambulance |
| INACTION | pants |
| ANIMAL? | heron |
| ANIMAL? | lamb |
| INACTION | panda |
| ANIMAL? | husky |
| INACTION | piranha |
| ANIMAL? | directory |
| ANIMAL? | cap |
| ANIMAL? | monkey |
| ANIMAL? | flask |
| INACTION | duck |
| ANIMAL? | rake |
| ANIMAL? | reptile |
| ANIMAL? | fish |
| ANIMAL? | stork |
| ANIMAL? | container |
| ANIMAL? | pencil |
| ANIMAL? | iguana |
| ANIMAL? | chisel |
| INACTION | raptor |
| INACTION | plate |
| ANIMAL? | shrimp |
| ANIMAL? | washing-machine |
| INACTION | syringe |
| ANIMAL? | lotto |
| ANIMAL? | sink |
| ANIMAL? | coat |
| INACTION | pillow |
| ANIMAL? | iceberg |
| INACTION | plow |
| INACTION | magnolia |
| ANIMAL? | perch |
| ANIMAL? | fortress |
| ANIMAL? | seagull |
| INACTION | weasel |
| INACTION | paddle |
| ANIMAL? | locker |
| ANIMAL? | mice |
| ANIMAL? | lion |
| INACTION | knife |
| INACTION | billy |
| ANIMAL? | bracelet |
| INACTION | spider |
| ANIMAL? | bell |
| INACTION | sheep |
| ANIMAL? | shark |
| INACTION | radio |
| ANIMAL? | gun |
| ANIMAL? | quail |
| INACTION | trout |
| ANIMAL? | notice |
| INACTION | mammoth |
| ANIMAL? | couch |
| INACTION | opossum |
| ANIMAL? | boat |
| INACTION | chimpanzee |
| INACTION | crawfish |
| ANIMAL? | cock |
| ANIMAL? | butterfly |
| INACTION | carnivore |
| ANIMAL? | seal |
| ANIMAL? | jenny |
| ANIMAL? | album |
| INACTION | gazelle |
| ANIMAL? | horse |
| ANIMAL? | moose |
| ANIMAL? | toad |
| ANIMAL? | racer |
| INACTION | train |
| ANIMAL? | revolver |
| INACTION | insect |
| ANIMAL? | ovals |
| INACTION | coffee pot |
| ANIMAL? | rhinoceros |
| ANIMAL? | eel |
| ANIMAL? | grasshopper |
| INACTION | kitty |
| INACTION | crutch |
| ANIMAL? | piano |
| ANIMAL? | baggage |
| ANIMAL? | tank |
| INACTION | crayfish |
| ANIMAL? | cheetah |
| ANIMAL? | penguin |
| ANIMAL? | boar |
| ANIMAL? | pony |
| ANIMAL? | blackbird |
| INACTION | partridge |
| INACTION | bison |
| INACTION | bear |
| ANIMAL? | ashtray |
| ANIMAL? | washbasin |
| ANIMAL? | mare |
| ANIMAL? | soap |
| ANIMAL? | ring |
| INACTION | locust |
| INACTION | rectangle |
| ANIMAL? | turkey |
| ANIMAL? | accordion |
| ANIMAL? | clarinet |
| ANIMAL? | plaster |
| ANIMAL? | sponge |
| ANIMAL? | bull |
| INACTION | mammal |
| INACTION | robin |
| ANIMAL? | tiger |
| INACTION | clinic |
| ANIMAL? | fence |
| ANIMAL? | microphone |
| ANIMAL? | camping |
| INACTION | shovel |
| INACTION | target |
| ANIMAL? | veal |
| ANIMAL? | bowl |
| ANIMAL? | broom |
| ANIMAL? | headband |
| INACTION | nightingale |
| ANIMAL? | hamster |
| ANIMAL? | axe |
| INACTION | accommodation |
| ANIMAL? | case |
| ANIMAL? | bump |
| INACTION | dome |
| ANIMAL? | doe |
| ANIMAL? | feline |
| ANIMAL? | vulture |
| ANIMAL? | cork |
| ANIMAL? | scorpion |
| INACTION | lamppost |
| ANIMAL? | chameleon |
| ANIMAL? | umbrella |
| INACTION | caterpillar |
| ANIMAL? | rifle |
| INACTION | cover |
| ANIMAL? | Dog |
| INACTION | checkbook |
| ANIMAL? | spool |
| ANIMAL? | frog |
| INACTION | fly |
| ANIMAL? | sweater |
| ANIMAL? | glue |
| ANIMAL? | flute |
| ANIMAL? | bottle |
| ANIMAL? | caribou |
| ANIMAL? | canapy |
| ANIMAL? | grizzly |
| INACTION | piston |
| INACTION | squirrel |
| ANIMAL? | ox |
| ANIMAL? | string |
| ANIMAL? | budgie |
| ANIMAL? | cow |
| INACTION | slipper |
| ANIMAL? | casserole |
| INACTION | bird |
| ANIMAL? | cauldron |
| INACTION | salmon |
| INACTION | sardine |
| INACTION | racket |
| ANIMAL? | dolphin |
| ANIMAL? | ferret |
| INACTION | beluga |
| INACTION | shearer |
| ANIMAL? | dalmatian |
| ANIMAL? | koala |
| ANIMAL? | dagger |
| ANIMAL? | owl |
| INACTION | ewe |
| ANIMAL? | hare |
| ANIMAL? | squid |
| ANIMAL? | cricket |
| ANIMAL? | gorilla |
| ANIMAL? | coyote |
| ANIMAL? | stopwatch |
| ANIMAL? | ostrich |
| INACTION | quadrate |
| ANIMAL? | stool |
| INACTION | crow |
| INACTION | glove |
| INACTION | dormitory |
| ANIMAL? | parcel |
| ANIMAL? | snail |
| INACTION | canary |
| ANIMAL? | rat |
| ANIMAL? | stagecoach |
| ANIMAL? | raven |
| ANIMAL? | goat |
| ANIMAL? | avalanche |
| ANIMAL? | polish |
| ANIMAL? | skate |
| ANIMAL? | drawer |
| ANIMAL? | donkey |
| ANIMAL? | dryer |
| ANIMAL? | trash |
| ANIMAL? | faucet |
| INACTION | mole |
| ANIMAL? | peaches |
| ANIMAL? | cannon |
| ANIMAL? | goose |
| ANIMAL? | skunk |
| ANIMAL? | petticoat |
| ANIMAL? | pike |
| INACTION | tire |
| ANIMAL? | computer |
| ANIMAL? | archive |
| ANIMAL? | roller |
| ANIMAL? | compass |
| INACTION | poodle |
| INACTION | review |
| INACTION | vase |
| INACTION | cupboard |
| ANIMAL? | castle |
| ANIMAL? | foal |
| ANIMAL? | pen |
| ANIMAL? | beret |
| ANIMAL? | snake |
| INACTION | chamois |
| INACTION | pelican |
| INACTION | beaver |
| ANIMAL? | oyster |
| INACTION | dynamite |
| ANIMAL? | stallion |
| ANIMAL? | screwdriver |
| ANIMAL? | lark |
| ANIMAL? | dove |
| INACTION | jewel |
| ANIMAL? | flag |
| ANIMAL? | ape |
| ANIMAL? | post |
| INACTION | pig |
| INACTION | ticket |
| INACTION | propeller |
| ANIMAL? | saw |
| ANIMAL? | dinosaur |
| ANIMAL? | bee |
| ANIMAL? | beetle |
| INACTION | giraffe |
| ANIMAL? | pyramid |
| ANIMAL? | jar |
| INACTION | fawn |
| INACTION | fork |
| ANIMAL? | tissue |
| ANIMAL? | herbivore |
| INACTION | chest |
| ANIMAL? | crocodile |
| INACTION | kettle |
| INACTION | elephant |
| ANIMAL? | cougar |
| ANIMAL? | ball |
| ANIMAL? | fox |
| ANIMAL? | cobra |
| ANIMAL? | aphid |
| ANIMAL? | hyena |
| ANIMAL? | buffalo |
| INACTION | mosquito |
| ANIMAL? | falcon |
| ANIMAL? | lama |
| ANIMAL? | calendar |
| ANIMAL? | label |
| INACTION | pool |
| INACTION | crab |
| INACTION | sparrow |
| INACTION | peacock |
| INACTION | cassette |
| INACTION | osprey |
| ANIMAL? | dragonfly |
| INACTION | lobster |
| ANIMAL? | hippopotamus |
| ANIMAL? | labrador |
| ANIMAL? | wolf |
| ANIMAL? | passport |
| ANIMAL? | dish |
| ANIMAL? | violin |
| ANIMAL? | shoe |
| ANIMAL? | garbage |
| ANIMAL? | swan |
| ANIMAL? | hat |
| INACTION | chicken |
| INACTION | key |
| ANIMAL? | tuna |
| ANIMAL? | cat |
| INACTION | jackal |
| ANIMAL? | rodent |
| ANIMAL? | mirror |
| ANIMAL? | comb |
| INACTION | circuit |
| INACTION | ant |
| ANIMAL? | seesaw |
| ANIMAL? | python |
| INACTION | kangaroo |
| INACTION | plane |
| INACTION | towel |
| INACTION | drill |
| ANIMAL? | squab |
| INACTION | padlock |
| ANIMAL? | fertilizer |
| ANIMAL? | sprinkler |
| ANIMAL? | pane |
| INACTION | hoop |
| ANIMAL? | turtle |
| ANIMAL? | lynx |
| ANIMAL? | airport |
| ANIMAL? | zebra |
| ANIMAL? | otter |
| INACTION | van |
| INACTION | ram |
| ANIMAL? | flake |
| ANIMAL? | stapler |
| ANIMAL? | shirt |
| ANIMAL? | cave |
| ANIMAL? | mummy |
| INACTION | necklace |
| ANIMAL? | ladybug |
| INACTION | sow |

### Table S13. List of the stimuli of the social role task.

|  |  |  | **Advantageousness** (favorability) 1-extremely unfavorable; 9-extremely favorable | | | **Ordinariness** 1-extremely ordinary; 9-extremely extraordinary | | | **Valence** 1-extremely pleasant; 9-extremely unpleasant | | | **Arousal** 1-extremely aroused; 9-extremely calm | | |
| --- | --- | --- | --- | --- | --- | --- | --- | --- | --- | --- | --- | --- | --- | --- |
| Predetermined Category |  | Frequency of occurrence (log) | Mean | SD | Median | Mean | SD | Median | Mean | SD | Median | Mean | SD | Median |
| Extraordinary favorable | Jesus | -2.09 | 5.78 | 2.5 | 5 | 7.28 | 2.11 | 8 | 3.94 | 2.38 | 4 | 5.72 | 2.73 | 5.5 |
| Extraordinary favorable | Harry Potter | -4.68 | 7 | 2.21 | 8 | 6.74 | 2.66 | 8 | 3.13 | 2.35 | 2 | 4.03 | 2.46 | 4 |
| Extraordinary favorable | knight | -3.34 | 5.72 | 2.37 | 6 | 6.17 | 2.38 | 7 | 4.1 | 2.09 | 5 | 4.1 | 2.38 | 4 |
| Extraordinary favorable | Buddha | -2.94 | 7.17 | 1.68 | 8 | 6.87 | 2.56 | 8 | 3.37 | 2.19 | 3 | 6.2 | 2.86 | 7.5 |
| Extraordinary favorable | ghostbuster | -6.58 | 5.03 | 2.37 | 5 | 6.74 | 2.65 | 8 | 4.58 | 2.14 | 4 | 4.32 | 2.29 | 4 |
| Extraordinary favorable | samurai | -3.89 | 5.39 | 2.17 | 5 | 6.61 | 2.12 | 7 | 5.03 | 2.02 | 5 | 4.42 | 2.29 | 5 |
| Extraordinary favorable | Peter Pan | -4.39 | 6.63 | 1.86 | 7 | 7.16 | 2.17 | 8 | 3.56 | 2.06 | 4 | 5.03 | 2.39 | 5 |
| Extraordinary favorable | Superman | -4.03 | 7.32 | 2.04 | 8 | 6.97 | 2.73 | 8 | 2.9 | 1.85 | 3 | 4 | 1.98 | 4 |
| Extraordinary favorable | fairy | -3.26 | 6.88 | 2.16 | 7 | 7.21 | 2.23 | 8 | 3.27 | 2.44 | 3 | 4.88 | 2.18 | 5 |
| Extraordinary favorable | Hindu God | -5.86 | 6.03 | 2.24 | 6 | 7.06 | 2.4 | 8 | 4.24 | 2.24 | 5 | 5.64 | 2.43 | 5 |
| Extraordinary favorable | prophet | -3.09 | 5.91 | 2.54 | 6 | 6.97 | 2.04 | 7 | 4.44 | 2.33 | 4.5 | 4.94 | 2.17 | 4.5 |
| Extraordinary favorable | Zeus | -3.5 | 5.53 | 2.16 | 5 | 6.93 | 2.49 | 8 | 4.27 | 2.23 | 5 | 4.53 | 1.91 | 4 |
| Extraordinary favorable | Einstein | -3.16 | 7.7 | 1.85 | 8 | 7.94 | 1.85 | 9 | 2.97 | 2.13 | 2 | 4.42 | 2.84 | 4 |
| Extraordinary favorable | mindreader | -6.22 | 4.66 | 2.77 | 5 | 6.59 | 2.41 | 8 | 5.62 | 2.38 | 5 | 5.14 | 2.46 | 5 |
| Extraordinary favorable | medieval king | -5.87 | 5.27 | 2.55 | 5 | 6.12 | 2.3 | 7 | 4.73 | 2.45 | 5 | 3.94 | 1.98 | 4 |
| Extraordinary favorable | God | -1.51 | 5.77 | 3.11 | 7 | 7.67 | 2.29 | 9 | 4.37 | 2.59 | 5 | 5.03 | 3.18 | 4 |
| Extraordinary favorable | mermaid | -4.38 | 5.91 | 2.18 | 6 | 7.38 | 2.03 | 8 | 3.13 | 2.32 | 2 | 4.25 | 1.88 | 4 |
| Extraordinary favorable | Joan of Arc | -4.21 | 5.69 | 2.19 | 5.5 | 6.47 | 2.17 | 7 | 4.41 | 2.33 | 4.5 | 4.91 | 1.91 | 5 |
| Extraordinary favorable | angel | -3.09 | 6.74 | 2.28 | 7 | 7.61 | 1.75 | 8 | 3.58 | 2.46 | 3 | 5.32 | 2.7 | 5 |
| Extraordinary favorable | elf | -4.25 | 5.88 | 1.77 | 5 | 7.69 | 1.69 | 8 | 3.47 | 2.11 | 3 | 5.16 | 2.53 | 5 |
| Extraordinary favorable | Gandhi | -2.95 | 7.09 | 2.43 | 8 | 7.13 | 2.28 | 8 | 2.84 | 2.54 | 2 | 5.13 | 2.97 | 5 |
| Extraordinary favorable | Noah | -3.37 | 6 | 2.08 | 6 | 6.78 | 2.21 | 7.5 | 4.38 | 2.18 | 5 | 5.5 | 2.02 | 6 |
| Extraordinary favorable | Napoleon | -3.01 | 4.45 | 2.26 | 5 | 6.72 | 2.46 | 8 | 5.41 | 2.34 | 5 | 4.86 | 2.49 | 5 |
| Extraordinary favorable | Robin Hood | -4.12 | 6.32 | 1.92 | 7 | 6.81 | 1.94 | 7 | 3.13 | 1.61 | 3 | 4.32 | 1.78 | 4 |
| Extraordinary favorable | Dalai Llama | -7.19 | 7.09 | 1.92 | 7.5 | 7.25 | 2 | 8 | 2.72 | 2.22 | 2 | 5.97 | 2.46 | 5.5 |
| Extraordinary favorable | Shakespeare | -2.71 | 6.65 | 2.04 | 7 | 7.45 | 1.98 | 8 | 3.19 | 2.09 | 3 | 5.03 | 2.34 | 5 |
| Extraordinary favorable | Hercules | -3.64 | 6.97 | 1.91 | 7 | 7.48 | 2.05 | 8 | 3.84 | 2.08 | 4 | 4.16 | 2.16 | 4 |
| Extraordinary favorable | Ironman | -5.11 | 6.55 | 2.4 | 7 | 6.85 | 2.49 | 8 | 3.76 | 2.36 | 4 | 3.42 | 2.12 | 3 |
| Extraordinary favorable | Santa Claus | -4.08 | 6.68 | 1.99 | 7 | 7.13 | 2.33 | 8 | 2.87 | 1.98 | 2 | 5.52 | 2.55 | 5 |
| Extraordinary favorable | Sigmund Freud | -3.88 | 5.55 | 2.05 | 5 | 6.52 | 1.91 | 7 | 4.19 | 2.01 | 4 | 4.52 | 1.96 | 4 |
| Extraordinary favorable | Batman | -4.24 | 6.93 | 2.38 | 8 | 7.3 | 2.35 | 8 | 3.77 | 2.65 | 3 | 3.4 | 2.22 | 3 |
| Extraordinary favorable | Brad Pitt | -5.32 | 6.56 | 2.11 | 7 | 5.38 | 2.81 | 6.5 | 3.53 | 2.54 | 3 | 4.53 | 2.41 | 4 |
| Extraordinary favorable | Angelina Jolie | -5.85 | 6.03 | 1.98 | 6 | 6 | 2.37 | 7 | 3.42 | 2.06 | 3 | 4.52 | 2.41 | 4 |
| Extraordinary favorable | Stephen Hawking | -5.05 | 7.03 | 1.92 | 8 | 7.16 | 2.11 | 8 | 3.23 | 1.61 | 3 | 5 | 2.53 | 5 |
| Extraordinary favorable | Madonna | -3.56 | 5.79 | 2.04 | 6 | 5.91 | 2.39 | 7 | 4.3 | 2.17 | 4 | 4.82 | 2.27 | 5 |
| Extraordinary favorable | Olympic athlete | -5.53 | 6.81 | 2.05 | 7 | 6.03 | 2.42 | 7 | 3.63 | 2.12 | 3 | 3.88 | 2.15 | 4 |
| Extraordinary favorable | Barack Obama | -4.47 | 6.69 | 2.29 | 7 | 6.03 | 2.76 | 7 | 4 | 2.78 | 3.5 | 4.41 | 2.54 | 4 |
| Extraordinary favorable | Bill Gates | -4.53 | 6.64 | 2.34 | 8 | 6.85 | 2.41 | 8 | 3.58 | 2.53 | 3 | 5.3 | 2.58 | 5 |
| Extraordinary favorable | Hillary Clinton | -4.75 | 6 | 2.35 | 6 | 5.67 | 2.39 | 6 | 4.15 | 2.31 | 4 | 5.18 | 2.14 | 5 |
| Extraordinary favorable | Moses | -2.82 | 6.52 | 1.9 | 6 | 6.81 | 2.37 | 7 | 3.9 | 1.99 | 4 | 5.19 | 2.33 | 5 |
| Extraordinary favorable | Pokemon trainer | -7.38 | 5.77 | 1.92 | 5.5 | 6.6 | 2.03 | 7 | 3.93 | 1.98 | 4 | 4.83 | 2.15 | 4.5 |
| Extraordinary favorable | Oprah Winfrey | -4.76 | 5.91 | 2.25 | 6 | 5.81 | 2.48 | 6 | 4.47 | 2.84 | 4 | 5.56 | 2.08 | 5 |
| Extraordinary favorable | Aladdin | -4.3 | 6.09 | 1.77 | 6 | 6.25 | 2.71 | 7 | 3.31 | 1.86 | 3 | 5.06 | 2.08 | 5 |
| Extraordinary favorable | Bono U2 | -4.32 | 5.61 | 2.16 | 6 | 5.9 | 2.3 | 6 | 4.32 | 2.36 | 4 | 4.65 | 2.03 | 5 |
| Extraordinary favorable | Winston Churchill | -3.87 | 6.25 | 1.78 | 6 | 6.19 | 1.79 | 6 | 4.41 | 1.92 | 5 | 4.91 | 1.92 | 5 |
| Extraordinary favorable | Jay-Z | -6.83 | 6.09 | 2.18 | 6.5 | 6.13 | 2.24 | 6.5 | 4.16 | 2.02 | 4 | 5.16 | 2.52 | 5 |
| Extraordinary favorable | Kate Middleton | -7.25 | 6.35 | 2.04 | 6 | 5.16 | 2.33 | 5 | 3.52 | 2.23 | 3 | 6.03 | 2.47 | 7 |
| Extraordinary favorable | Prince William | -4.43 | 5.6 | 1.9 | 5 | 5.27 | 2.33 | 6 | 4.5 | 2.26 | 5 | 5.9 | 2.23 | 6 |
| Extraordinary favorable | Serena Williams | -6 | 6.09 | 2.04 | 6 | 6.39 | 2.19 | 7 | 3.91 | 2.11 | 4 | 4.91 | 2.02 | 5 |
| Extraordinary favorable | Queen Elizabeth II | -4.62 | 5.44 | 2.15 | 5 | 6.44 | 2.31 | 7 | 4.03 | 2.32 | 4 | 5.66 | 2.31 | 5 |
| Extraordinary favorable | Steve Jobs | -5.04 | 6.57 | 2.13 | 7.5 | 6.83 | 2.42 | 8 | 4.3 | 2.59 | 4 | 5.17 | 1.93 | 5 |
| Extraordinary favorable | Pierre Trudeau | -4.87 | 5.74 | 2.14 | 6 | 6.13 | 2.13 | 6 | 4.58 | 1.93 | 5 | 5.16 | 2.07 | 5 |
| Extraordinary favorable | Bob Marley | -4.86 | 6.94 | 1.41 | 7 | 6.61 | 1.98 | 7 | 3.16 | 1.71 | 3 | 5.68 | 2.43 | 6 |
| Extraordinary favorable | Spiderman | -5.24 | 7.06 | 1.87 | 8 | 7.76 | 1.66 | 8 | 3.45 | 2.2 | 3 | 4.06 | 2.26 | 3 |
| Extraordinary favorable | Che Guevara | -4.55 | 5.39 | 1.96 | 5 | 6.39 | 2.25 | 7 | 4.84 | 1.93 | 5 | 3.94 | 1.79 | 5 |
| Extraordinary favorable | Mark Zuckerberg | -7.23 | 5.58 | 2.2 | 6 | 6.19 | 1.96 | 7 | 4.9 | 1.97 | 5 | 5.23 | 1.84 | 5 |
| Extraordinary favorable | Charles Darwin | -4.1 | 6.91 | 2.12 | 7 | 7.09 | 2.01 | 8 | 3.41 | 2.09 | 3 | 4.84 | 2.17 | 5 |
| Extraordinary favorable | Gulliver | -4.02 | 5.68 | 1.45 | 5 | 5.97 | 1.97 | 6 | 4.58 | 1.75 | 5 | 5.52 | 1.86 | 5 |
| Extraordinary favorable | Cinderella | -3.92 | 6.56 | 2.17 | 7 | 6.41 | 2.59 | 7 | 3.44 | 2.26 | 2.5 | 5.41 | 2.2 | 5 |
| Extraordinary favorable | Marilyn Monroe | -4.35 | 5.94 | 2.01 | 7 | 6.12 | 2.26 | 7 | 3.76 | 1.68 | 4 | 4.94 | 2.12 | 5 |
| Extraordinary favorable | Princess Diana | -4.86 | 6.06 | 2.21 | 6 | 5.81 | 2.3 | 6 | 3.65 | 2.46 | 3 | 5.58 | 2.13 | 5 |
| Extraordinary favorable | Nelson Mandela | -4.38 | 7.31 | 1.75 | 7.5 | 6.63 | 2.51 | 7.5 | 2.75 | 1.8 | 2 | 4.59 | 2.24 | 5 |
| Extraordinary favorable | Cristiano†Ronaldo | -7.43 | 6.55 | 1.79 | 6 | 6.26 | 2.21 | 7 | 3.52 | 1.96 | 4 | 4.16 | 2.03 | 4 |
| Extraordinary favorable | Michael Phelps | -6.5 | 6.35 | 2.07 | 7 | 6.94 | 2.46 | 8 | 4.03 | 2.27 | 4 | 4.42 | 2.31 | 4 |
| Extraordinary favorable | Zorro | -4.82 | 6.06 | 1.82 | 6 | 6.79 | 2.16 | 7 | 4.39 | 2 | 4 | 4.73 | 2.27 | 5 |
| Extraordinary favorable | Salvador Dali | -4.75 | 6.5 | 2.08 | 6.5 | 6.4 | 2.24 | 7 | 3.43 | 2.14 | 4 | 4.37 | 2.2 | 5 |
| Extraordinary favorable | Cupid | -4.01 | 5.82 | 2.3 | 6 | 6.82 | 2.4 | 8 | 3.82 | 2.17 | 3 | 5.06 | 2.29 | 5 |
| Extraordinary favorable | Bugs Bunny | -5.01 | 6.65 | 1.6 | 7 | 6.26 | 2.52 | 7 | 3.19 | 1.85 | 3 | 4.97 | 1.94 | 5 |
| Extraordinary favorable | Uncle Sam | -4.15 | 5 | 2.37 | 5 | 5.19 | 2.56 | 6 | 4.74 | 2.45 | 5 | 5.65 | 2.06 | 5 |
| Extraordinary favorable | wizard | -3.71 | 6.37 | 2.22 | 6.5 | 7.3 | 2.39 | 8 | 4.03 | 2.16 | 4.5 | 4.33 | 2.37 | 5 |
| Extraordinary favorable | Three Wise Men | -4.38 | 5.74 | 1.75 | 5 | 5.58 | 2.25 | 6 | 4.26 | 2.24 | 5 | 5.77 | 2.23 | 5 |
| Extraordinary favorable | Pied Piper | -4.78 | 5.48 | 1.42 | 5 | 5.88 | 2.06 | 5 | 4.27 | 1.33 | 5 | 5.82 | 1.98 | 5 |
| Extraordinary favorable | army general | -5.03 | 4.7 | 2.45 | 5 | 5.17 | 1.93 | 5 | 5.53 | 2.39 | 6 | 4.97 | 2.33 | 5 |
| Extraordinary favorable | FBI agent | -4.5 | 5.63 | 2.31 | 6 | 6.03 | 1.58 | 6 | 5.13 | 2.41 | 5 | 4.19 | 2.15 | 4 |
| Extraordinary favorable | sultan | -3.74 | 5.26 | 2.18 | 5 | 5.68 | 2.34 | 5 | 4.84 | 2.16 | 5 | 5.35 | 2.56 | 5 |
| Extraordinary favorable | ninja | -4.92 | 5.66 | 2.12 | 6 | 6.53 | 1.98 | 7 | 4.34 | 2.18 | 4 | 3.94 | 2.05 | 4 |
| Extraordinary favorable | Native Indian | -5.52 | 5.9 | 1.86 | 5 | 5.2 | 2.02 | 5 | 4.37 | 1.63 | 5 | 5.3 | 2.12 | 5 |
| Extraordinary favorable | tightrope walker | -5.28 | 5.58 | 1.8 | 5 | 6.35 | 2.06 | 7 | 4.48 | 2.13 | 4 | 4.13 | 2.32 | 3 |
| Extraordinary favorable | Pharaoh | -4.78 | 5.79 | 2.19 | 6 | 6.72 | 2.23 | 8 | 5 | 2.36 | 5 | 5.14 | 2.08 | 5 |
| Extraordinary favorable | satyr | -4.48 | 4.94 | 1.69 | 5 | 6.24 | 2.02 | 6 | 5 | 1.41 | 5 | 4.85 | 1.86 | 5 |
| Extraordinary favorable | faun | -4.94 | 6.19 | 1.23 | 6 | 5.28 | 2.3 | 5 | 3.84 | 2 | 4 | 5.75 | 1.83 | 6 |
| Extraordinary favorable | leprechaun | -5.03 | 5.27 | 2.52 | 5 | 7.4 | 2.01 | 8 | 4.87 | 2.22 | 5 | 4.33 | 2.02 | 4 |
| Extraordinary favorable | psychic | -3.09 | 4.5 | 2.17 | 5 | 6.25 | 2.23 | 6 | 5.44 | 2.33 | 5 | 4.53 | 2.2 | 4 |
| Extraordinary favorable | Alice in Wonderland | -4.49 | 6.06 | 2.03 | 6.5 | 6.63 | 2.32 | 7.5 | 3.59 | 1.88 | 4 | 4.81 | 2.25 | 4.5 |
| Extraordinary unfavorable | caveman | -4.96 | 4.39 | 1.86 | 5 | 4.84 | 2.58 | 5 | 5.58 | 2 | 5 | 4.61 | 2.04 | 5 |
| Extraordinary unfavorable | Invisible Man | -4.49 | 5.13 | 2.13 | 5 | 7.9 | 1.73 | 9 | 4.8 | 1.86 | 5 | 4.67 | 2.25 | 5 |
| Extraordinary unfavorable | werewolf | -4.53 | 3.25 | 2.11 | 2 | 6.91 | 2.51 | 8 | 6.56 | 2.2 | 7 | 3.5 | 2.23 | 3 |
| Extraordinary unfavorable | evil wizard | -6.03 | 3 | 2.46 | 2 | 7.25 | 2.38 | 8 | 7.13 | 2.21 | 8 | 4.03 | 2.52 | 3 |
| Extraordinary unfavorable | alien | -2.89 | 4.63 | 2.45 | 5 | 7.34 | 2.25 | 8 | 6.03 | 2.22 | 6 | 3.75 | 2.33 | 4 |
| Extraordinary unfavorable | centaur | -4.6 | 5.52 | 1.96 | 5 | 7.03 | 2.12 | 8 | 4.58 | 2.25 | 5 | 4.74 | 1.86 | 5 |
| Extraordinary unfavorable | Elephant Man | -5.23 | 3.78 | 2.68 | 3 | 6.97 | 2.36 | 8 | 5.97 | 2.86 | 6.5 | 4.28 | 2.39 | 4.5 |
| Extraordinary unfavorable | Frankenstein | -3.84 | 4 | 2.7 | 4 | 7.26 | 1.93 | 8 | 6.32 | 2.51 | 7 | 4.55 | 2.55 | 5 |
| Extraordinary unfavorable | Hades | -4 | 3.25 | 2.44 | 2.5 | 7 | 2.3 | 8 | 6.63 | 2.42 | 7 | 3.97 | 2.38 | 4 |
| Extraordinary unfavorable | Devil | -3.28 | 3.09 | 2.79 | 1.5 | 7.09 | 2.26 | 8 | 7.69 | 2.12 | 9 | 3.63 | 2.5 | 3 |
| Extraordinary unfavorable | mummy | -4.01 | 4.81 | 2.77 | 5 | 6.74 | 2.31 | 8 | 5.74 | 2.85 | 7 | 4.19 | 2.39 | 3 |
| Extraordinary unfavorable | hunchback | -4.56 | 3.69 | 2.12 | 3.5 | 5.53 | 2.2 | 6 | 6.28 | 2.16 | 7 | 5.31 | 1.96 | 5 |
| Extraordinary unfavorable | Captain Hook | -5.4 | 4.04 | 2.08 | 4 | 6.54 | 2.28 | 7 | 5.25 | 2.27 | 5 | 4.39 | 1.93 | 4 |
| Extraordinary unfavorable | slave | -2.61 | 1.91 | 1.78 | 1 | 5.34 | 2.61 | 6 | 8.19 | 1.35 | 9 | 3.72 | 2.04 | 4 |
| Extraordinary unfavorable | dwarf | -3.5 | 4.44 | 1.85 | 5 | 5.91 | 1.86 | 6 | 4.63 | 1.54 | 5 | 6 | 1.41 | 5 |
| Extraordinary unfavorable | ghost | -3.12 | 3.44 | 2.31 | 3 | 7.5 | 2.02 | 8 | 6.47 | 2.17 | 7 | 3.78 | 2.15 | 4 |
| Extraordinary unfavorable | Dr. Jekyll | -5.38 | 4.61 | 2.17 | 5 | 6.29 | 2.16 | 7 | 5.32 | 2.27 | 5 | 5.19 | 2.29 | 5 |
| Extraordinary unfavorable | evil clown | -6.81 | 2.74 | 2.25 | 2 | 6.23 | 2.38 | 7 | 7.29 | 2.24 | 8 | 3.55 | 2.08 | 3 |
| Extraordinary unfavorable | jester | -4.43 | 5.58 | 1.58 | 6 | 5.58 | 2.08 | 6 | 4.39 | 1.66 | 4 | 4.73 | 1.97 | 5 |
| Extraordinary unfavorable | vampire | -3.83 | 3.34 | 2.16 | 3 | 7.31 | 2.04 | 8 | 6.44 | 2.63 | 7 | 3.56 | 2.09 | 3 |
| Extraordinary unfavorable | Cyclops | -4.36 | 4.45 | 2.39 | 5 | 7.24 | 2.18 | 8 | 6.09 | 2.47 | 7 | 4.33 | 2.13 | 5 |
| Extraordinary unfavorable | The Joker | -4.67 | 3.83 | 2.57 | 3 | 6.97 | 2.5 | 8 | 6.13 | 2.75 | 6.5 | 3.6 | 2.14 | 3 |
| Extraordinary unfavorable | Medusa | -4.15 | 3.48 | 2.77 | 2 | 7.26 | 2.41 | 8 | 6.35 | 2.83 | 8 | 3.74 | 2.16 | 4 |
| Extraordinary unfavorable | Hitler | -2.73 | 2.38 | 2.42 | 1 | 7.66 | 1.98 | 8.5 | 7.97 | 1.94 | 9 | 2.78 | 2.43 | 1.5 |
| Extraordinary unfavorable | alien abductee | -7.12 | 3.32 | 2.36 | 2 | 8 | 1.81 | 9 | 6.39 | 2.14 | 7 | 3.48 | 2.74 | 3 |
| Extraordinary unfavorable | gladiator | -4.66 | 4.9 | 2.12 | 5 | 6.45 | 2.16 | 7 | 6.1 | 2.04 | 6 | 3.77 | 2.23 | 4 |
| Extraordinary unfavorable | Grim Reaper | -5.24 | 3 | 2.58 | 2 | 6.39 | 2.93 | 8 | 6.55 | 2.94 | 8 | 3.84 | 2.5 | 3 |
| Extraordinary unfavorable | cyborg | -4.58 | 4.83 | 1.98 | 5 | 5.97 | 2.34 | 7 | 4.9 | 1.99 | 5 | 4.47 | 2.1 | 5 |
| Extraordinary unfavorable | zombie | -4.41 | 2.6 | 2.11 | 1.5 | 7.53 | 2.42 | 8.5 | 7.07 | 2.38 | 8 | 3.27 | 2.39 | 2 |
| Extraordinary unfavorable | pirate | -3.72 | 3.81 | 2.16 | 3.5 | 5.94 | 2.02 | 6 | 6 | 2.42 | 6 | 4.09 | 1.99 | 4 |
| Extraordinary unfavorable | Darth Vader | -5.07 | 3.88 | 2.57 | 3.5 | 6.44 | 3.04 | 8 | 6.56 | 2.21 | 7 | 4.09 | 2.63 | 3 |
| Extraordinary unfavorable | witch | -3.3 | 3.35 | 2.32 | 3 | 7.29 | 2.1 | 8 | 6.84 | 2.16 | 7 | 4.42 | 2.39 | 4 |
| Extraordinary unfavorable | Green Goblin | -6.32 | 3.71 | 2.19 | 4 | 7.03 | 2.24 | 8 | 6.35 | 2.12 | 7 | 4 | 1.71 | 4 |
| Extraordinary unfavorable | leper | -4.19 | 3.28 | 2.53 | 2 | 6.16 | 2 | 6 | 6.81 | 2.24 | 7 | 4.66 | 2.06 | 4.5 |
| Extraordinary unfavorable | Kim Jong II | -5.32 | 3.55 | 2.43 | 3 | 6.14 | 2.61 | 7 | 6.76 | 2.31 | 8 | 3.59 | 2.06 | 3 |
| Extraordinary unfavorable | Joseph Stalin | -4.62 | 3.29 | 2.55 | 2 | 6.68 | 2.45 | 7 | 6.74 | 2.46 | 8 | 3.39 | 2.32 | 3 |
| Extraordinary unfavorable | Bin Laden | -4.58 | 1.77 | 1.59 | 1 | 6.57 | 2.66 | 8 | 8.43 | 1.28 | 9 | 3.63 | 2.57 | 3 |
| Extraordinary unfavorable | Muammar Gaddafi | -6.02 | 3.06 | 2.14 | 2 | 5.91 | 2.52 | 6 | 6.91 | 2.15 | 8 | 3.91 | 2.16 | 4 |
| Extraordinary unfavorable | Hannibal Lecter | -5.58 | 2.84 | 2 | 2.5 | 6.44 | 2.66 | 7.5 | 7.13 | 1.93 | 7 | 3.94 | 2.55 | 3 |
| Extraordinary unfavorable | Cruella DeVil | -7.18 | 3.27 | 1.95 | 2.5 | 6.27 | 1.93 | 6.5 | 7.07 | 1.89 | 8 | 3.9 | 2.04 | 4 |
| Extraordinary unfavorable | Sumo Wrestler | -7.12 | 5.19 | 1.85 | 5 | 5.39 | 2.09 | 5 | 4.65 | 2.03 | 5 | 5 | 1.95 | 5 |
| Extraordinary unfavorable | Sword Swallower | -6.75 | 3.94 | 2.1 | 4 | 6.61 | 1.91 | 7 | 5.84 | 2.34 | 6 | 3.58 | 1.86 | 4 |
| Extraordinary unfavorable | ogre | -4.38 | 4.47 | 2.4 | 4.5 | 7.41 | 1.85 | 8 | 6.47 | 2.23 | 7 | 4.63 | 2.31 | 5 |
| Extraordinary unfavorable | WWF Wrestler | -7.19 | 4.63 | 2.04 | 5 | 5.53 | 2.01 | 6 | 5.17 | 2.21 | 5 | 3.93 | 2.48 | 4 |
| Extraordinary unfavorable | muscleman | -5.75 | 4.77 | 2.19 | 5 | 4.68 | 1.78 | 5 | 5.1 | 2.15 | 5 | 4.71 | 1.88 | 5 |
| Extraordinary unfavorable | conjoined twins | -5.39 | 3.97 | 2.56 | 4 | 7.38 | 2.24 | 8 | 6.22 | 2.71 | 6.5 | 4.25 | 2.03 | 4 |
| Extraordinary unfavorable | contortionist | -5.38 | 5.19 | 2.02 | 5 | 6 | 2.5 | 7 | 4.71 | 2.07 | 4 | 4.39 | 2.11 | 4 |
| Extraordinary unfavorable | fire eater | -6.28 | 4.26 | 2.29 | 4 | 6.61 | 2.25 | 7 | 5.81 | 2.34 | 6 | 3.52 | 2.11 | 3 |
| Extraordinary unfavorable | genocide victim | -7.56 | 1.91 | 1.87 | 1 | 5.91 | 2.53 | 6.5 | 7.47 | 2.58 | 9 | 2.94 | 1.98 | 3 |
| Extraordinary unfavorable | Ted Bundy | -5.23 | 3.34 | 2.3 | 2 | 5.9 | 2.51 | 5 | 6.69 | 2.32 | 7 | 3.55 | 1.99 | 5 |
| Extraordinary unfavorable | evil stepmother | -5.86 | 2.43 | 2.16 | 2 | 5.27 | 2.46 | 5 | 8.03 | 1.59 | 8.5 | 3.77 | 2.57 | 3 |
| Extraordinary unfavorable | seawitch | -7.35 | 4.06 | 2.69 | 4 | 6.94 | 2.27 | 8 | 6.28 | 2.25 | 6.5 | 4.16 | 2.49 | 4 |
| Ordinary unfavorable | beekeeper | -4.83 | 5.16 | 2.34 | 5 | 5.32 | 2.12 | 6 | 4.97 | 2.8 | 4 | 5.45 | 2.08 | 5 |
| Ordinary unfavorable | garbage man | -5.54 | 4.81 | 2.16 | 5 | 3.13 | 1.72 | 3 | 6.13 | 1.74 | 6 | 6.5 | 1.9 | 7 |
| Ordinary unfavorable | KKK member | -6.77 | 2.21 | 1.96 | 1 | 6.73 | 2.25 | 8 | 8.12 | 1.56 | 9 | 2.79 | 1.87 | 2 |
| Ordinary unfavorable | terrorist | -3.16 | 2.21 | 2.36 | 1 | 6.79 | 2.29 | 7 | 8.27 | 1.74 | 9 | 2.76 | 2.18 | 2 |
| Ordinary unfavorable | cripple | -3.96 | 3.53 | 2.5 | 3 | 5.09 | 2.33 | 5 | 6.84 | 2.1 | 7 | 5.03 | 2.18 | 5 |
| Ordinary unfavorable | tattooed man | -5.84 | 4.5 | 2.27 | 5 | 4 | 2.17 | 4 | 4.87 | 1.93 | 5 | 4.7 | 2.07 | 4.5 |
| Ordinary unfavorable | weed smoker | -7.62 | 4.59 | 2.45 | 5 | 3.25 | 2.02 | 3 | 4.75 | 2.58 | 5 | 5.75 | 2.58 | 6 |
| Ordinary unfavorable | convict | -3.59 | 2.69 | 2.24 | 2 | 4.97 | 2.26 | 5 | 7.25 | 1.98 | 8 | 3.94 | 2.09 | 4 |
| Ordinary unfavorable | abusive guard | -7.85 | 3.03 | 2.91 | 2 | 5.16 | 2.33 | 5 | 7.5 | 2.44 | 9 | 2.88 | 2.03 | 2 |
| Ordinary unfavorable | poor child | -4.5 | 2.79 | 2.06 | 2 | 3.48 | 1.91 | 3 | 7.88 | 1.56 | 8 | 4.09 | 1.79 | 4 |
| Ordinary unfavorable | homeless person | -6.08 | 3.03 | 2.13 | 2 | 3.48 | 2.14 | 3 | 7.52 | 1.54 | 8 | 4.67 | 2.34 | 4 |
| Ordinary unfavorable | burglar | -4.02 | 2.45 | 1.96 | 2 | 5.06 | 2.16 | 5 | 7.23 | 2.32 | 8 | 4.19 | 2.4 | 4 |
| Ordinary unfavorable | rioter | -5.42 | 3.66 | 1.94 | 3 | 4.56 | 1.81 | 5 | 5.94 | 2.2 | 7 | 3.94 | 2.15 | 3 |
| Ordinary unfavorable | rebels | -3.2 | 4.56 | 2.2 | 4.5 | 4.97 | 2.09 | 5 | 5.91 | 2.28 | 6 | 4.25 | 2.05 | 4 |
| Ordinary unfavorable | child soldier | -6.2 | 1.93 | 1.87 | 1 | 6.33 | 2.25 | 6.5 | 8.33 | 1.54 | 9 | 2.8 | 2.22 | 2 |
| Ordinary unfavorable | amputee | -4.66 | 3.45 | 2.31 | 3 | 5.7 | 2.11 | 6 | 6.33 | 2.03 | 7 | 4.82 | 1.57 | 5 |
| Ordinary unfavorable | anorexic | -4.38 | 2.59 | 2.31 | 1.5 | 4.84 | 2.36 | 5 | 7.81 | 1.75 | 8.5 | 3.69 | 2.22 | 3.5 |
| Ordinary unfavorable | goth | -5.16 | 3.41 | 1.56 | 4 | 4.56 | 2.2 | 5 | 6.31 | 1.77 | 6 | 5.47 | 2.02 | 5 |
| Ordinary unfavorable | dictator | -3.61 | 2.94 | 2.44 | 2 | 6.06 | 2.36 | 7 | 7.15 | 2.36 | 8 | 4.12 | 2.58 | 4 |
| Ordinary unfavorable | domestically abused | -7.51 | 1.65 | 1.31 | 1 | 4.35 | 2.12 | 4 | 8.1 | 1.97 | 9 | 2.68 | 1.87 | 2 |
| Ordinary unfavorable | computer nerd | -5.85 | 5.81 | 2.07 | 6 | 4.28 | 2.04 | 4 | 4.66 | 2.22 | 5 | 5.63 | 1.91 | 5.5 |
| Ordinary unfavorable | protestor | -5.36 | 4.7 | 2.31 | 5 | 4.55 | 2.03 | 5 | 5.48 | 2.43 | 5 | 3.94 | 2.26 | 4 |
| Ordinary unfavorable | obese person | -5.4 | 3.13 | 1.98 | 3 | 3.56 | 1.98 | 3 | 6.78 | 1.91 | 7 | 4.75 | 1.95 | 5 |
| Ordinary unfavorable | starving child | -5.65 | 2.85 | 2.8 | 1 | 4.52 | 2.37 | 5 | 7.97 | 1.93 | 9 | 4 | 2.49 | 4 |
| Ordinary unfavorable | pregnant teen | -5.97 | 3.06 | 2.2 | 2 | 4.28 | 1.99 | 4 | 6.63 | 2.2 | 7 | 4.69 | 2.05 | 4 |
| Ordinary unfavorable | suicidal person | -5.3 | 2.61 | 2.25 | 1 | 5.55 | 1.77 | 5 | 7.23 | 2.2 | 8 | 2.68 | 2.04 | 2 |
| Ordinary unfavorable | carjacker | -6.2 | 2.32 | 1.89 | 1 | 4.61 | 2.04 | 5 | 7.74 | 1.79 | 8 | 4.39 | 2.38 | 5 |
| Ordinary unfavorable | neo-nazi | -6.17 | 2.16 | 1.95 | 1 | 5.97 | 2.29 | 6 | 8.03 | 1.51 | 9 | 3.38 | 2.3 | 3 |
| Ordinary unfavorable | abused man | -6.42 | 2.29 | 1.97 | 2 | 4.9 | 2.43 | 5 | 8 | 1.44 | 8 | 3.68 | 1.92 | 3 |
| Ordinary unfavorable | executioner | -4.07 | 3.2 | 2.52 | 2 | 6.23 | 2.34 | 6.5 | 7.27 | 2.26 | 9 | 3.8 | 2.43 | 3 |
| Ordinary unfavorable | gangster | -4.01 | 3.1 | 1.73 | 3 | 4.43 | 1.85 | 5 | 6.8 | 1.86 | 7 | 4.2 | 2.14 | 4 |
| Ordinary unfavorable | self-immolator | -7.53 | 3.84 | 2.38 | 4 | 6.13 | 2.19 | 6 | 6 | 2.59 | 6 | 3.94 | 2.25 | 4 |
| Ordinary unfavorable | cleaning man | -6.23 | 5.63 | 1.85 | 5.5 | 3.13 | 2.27 | 2 | 4.23 | 2.03 | 5 | 6.83 | 1.66 | 7 |
| Ordinary unfavorable | emo | -4.98 | 3.27 | 1.84 | 3 | 4.88 | 2.58 | 5 | 6.48 | 2.12 | 7 | 5.55 | 2.37 | 6 |
| Ordinary unfavorable | grieving person | -5.54 | 3.35 | 2.18 | 3 | 3.77 | 1.91 | 4 | 7.45 | 1.36 | 8 | 5.42 | 2.16 | 5 |
| Ordinary unfavorable | drunk driver | -5.06 | 2.06 | 1.82 | 1 | 4.06 | 2.34 | 4 | 8.42 | 1.23 | 9 | 3.35 | 2.06 | 3 |
| Ordinary unfavorable | bullfighter | -4.9 | 4.16 | 2.38 | 4 | 5.81 | 1.91 | 6 | 6.47 | 2.02 | 7 | 3.69 | 2.07 | 3 |
| Ordinary unfavorable | burn victim | -4.75 | 1.93 | 1.56 | 1 | 5.45 | 2.29 | 5 | 8.24 | 1.38 | 9 | 3.21 | 1.93 | 3 |
| Ordinary unfavorable | Muslim extremists | -5.6 | 2.65 | 1.96 | 2 | 6.19 | 2.27 | 7 | 7.35 | 1.82 | 8 | 3.19 | 1.94 | 3 |
| Ordinary unfavorable | pickpocket | -4.81 | 2.81 | 2.23 | 2 | 4.84 | 2.4 | 5 | 7.68 | 1.8 | 8 | 4.03 | 2.01 | 4 |
| Ordinary unfavorable | punk | -3.92 | 4.23 | 1.91 | 4 | 4.23 | 2.03 | 5 | 5.74 | 1.86 | 6 | 5.06 | 1.93 | 5 |
| Ordinary unfavorable | abused child | -4.78 | 1.94 | 1.77 | 1 | 5.24 | 2.31 | 5 | 8.27 | 1.28 | 9 | 3.12 | 2.07 | 3 |
| Ordinary unfavorable | prostitute | -3.56 | 3.45 | 2.31 | 3 | 4.83 | 2.24 | 5 | 6.69 | 1.97 | 7 | 4.17 | 2.04 | 4 |
| Ordinary unfavorable | losing boxer | -7.39 | 3.34 | 2.06 | 3 | 3.94 | 2.03 | 4 | 6.81 | 1.77 | 7 | 4.88 | 2.04 | 5 |
| Ordinary unfavorable | heroin user | -5.73 | 2.34 | 2.04 | 1 | 5.06 | 2.85 | 5 | 8.03 | 1.51 | 9 | 4.06 | 2.42 | 3.5 |
| Ordinary unfavorable | smoker | -3.93 | 2.84 | 2.13 | 2 | 2.87 | 2.26 | 2 | 6.71 | 2.25 | 7 | 5.74 | 2.18 | 5 |
| Ordinary unfavorable | shoplifter | -5.1 | 3.31 | 2.33 | 2.5 | 4.09 | 2.43 | 3.5 | 6.59 | 2.54 | 7.5 | 4.63 | 2.04 | 4.5 |
| Ordinary unfavorable | alcoholic | -3.16 | 2.64 | 2.34 | 2 | 3.88 | 2.2 | 4 | 7.27 | 2.23 | 8 | 4.55 | 2.37 | 5 |
| Ordinary unfavorable | attacked woman | -7.17 | 1.9 | 1.47 | 1 | 4.68 | 2.43 | 5 | 8.55 | 0.93 | 9 | 3.48 | 2.53 | 2 |
| Ordinary unfavorable | terminal patient | -5.5 | 2.47 | 2.03 | 2 | 4.56 | 2.34 | 5 | 7.41 | 2.08 | 8 | 4.59 | 2.2 | 4 |
| Ordinary unfavorable | executed person | -6.39 | 3 | 2.7 | 1 | 5.56 | 2.42 | 6 | 7.41 | 2.17 | 8.5 | 3.5 | 2.29 | 3 |
| Ordinary unfavorable | gang member | -4.83 | 2.87 | 2.03 | 2.5 | 4.43 | 2.27 | 4 | 7.5 | 1.59 | 8 | 3.83 | 2.26 | 3 |
| Ordinary unfavorable | domestic abuser | -7.27 | 1.94 | 1.81 | 1 | 4.47 | 2.51 | 4 | 8.22 | 1.62 | 9 | 2.72 | 2.29 | 2 |
| Ordinary unfavorable | gambler | -3.91 | 3.44 | 2.29 | 3 | 3.63 | 2.11 | 3 | 6.34 | 2.04 | 7 | 5 | 1.9 | 5 |
| Ordinary unfavorable | transvestite | -4.47 | 4.97 | 2.06 | 5 | 5.53 | 1.98 | 6 | 4.63 | 1.97 | 5 | 4.67 | 2.12 | 5 |
| Ordinary unfavorable | pubescent teen | -7.74 | 4.1 | 1.96 | 4 | 2.52 | 2.05 | 2 | 6.65 | 1.58 | 6 | 5.13 | 2.47 | 5 |
| Ordinary unfavorable | blind | -2.65 | 2.8 | 1.95 | 2 | 5.73 | 2.21 | 6 | 7.27 | 1.84 | 8 | 5.13 | 1.96 | 5 |
| Ordinary unfavorable | bully | -3.79 | 2.3 | 1.78 | 2 | 3.52 | 2.15 | 3 | 7.58 | 1.8 | 8 | 3.76 | 2.24 | 3 |
| Ordinary unfavorable | child worker | -6 | 4.13 | 3.07 | 3 | 4.39 | 2.29 | 5 | 6.1 | 3.02 | 7 | 5 | 2.52 | 5 |
| Ordinary unfavorable | murder victim | -3.99 | 2.13 | 1.91 | 1 | 5.69 | 2.58 | 6 | 8.25 | 1.65 | 9 | 2.84 | 2.16 | 2 |
| Ordinary unfavorable | vandal | -5 | 3.83 | 1.91 | 4 | 4.33 | 2.01 | 4 | 6.33 | 2.01 | 6 | 4.43 | 1.98 | 4 |
| Ordinary unfavorable | chauffeur | -3.87 | 5.48 | 2.11 | 6 | 3.29 | 1.88 | 3 | 4.55 | 1.8 | 5 | 6.77 | 1.52 | 7 |
| Ordinary unfavorable | hillbilly | -4.65 | 4.1 | 1.84 | 4.5 | 4.03 | 1.96 | 4.5 | 5.47 | 1.94 | 6 | 5.7 | 2.02 | 5 |
| Ordinary unfavorable | plowman | -5.14 | 4.94 | 1.85 | 5 | 3.67 | 2.26 | 4 | 5.15 | 1.46 | 5 | 6.36 | 1.8 | 7 |
| Ordinary unfavorable | nomad | -4.29 | 5.13 | 2.46 | 5 | 5.23 | 2.12 | 5 | 4.81 | 2.21 | 5 | 5.61 | 1.91 | 5 |
| Ordinary unfavorable | window washer | -5.75 | 4.9 | 2.02 | 5 | 3 | 1.95 | 2 | 5.3 | 1.78 | 5 | 6.47 | 1.76 | 7 |
| Ordinary unfavorable | rapper | -4.87 | 5.52 | 2.16 | 6 | 5.03 | 2.08 | 6 | 4.45 | 1.9 | 5 | 4.24 | 1.57 | 5 |
| Ordinary unfavorable | showgirl | -4.15 | 4.84 | 2.25 | 5 | 4.35 | 2.03 | 4 | 4.48 | 2.08 | 5 | 4.68 | 2.04 | 5 |
| Ordinary unfavorable | plumber | -4.15 | 5.6 | 1.75 | 5.5 | 3.57 | 1.81 | 4 | 4.77 | 2.01 | 5 | 6.17 | 1.82 | 6 |
| Ordinary unfavorable | old man | -2.78 | 5.23 | 1.93 | 5 | 2.35 | 1.74 | 2 | 4.81 | 1.83 | 5 | 6.13 | 1.93 | 6 |
| Ordinary unfavorable | sick person | -4.32 | 3 | 2.25 | 2 | 2.62 | 1.74 | 2 | 7.76 | 1.72 | 8 | 5.45 | 1.82 | 5 |
| Ordinary unfavorable | snake charmer | -5.41 | 5.26 | 1.93 | 5 | 6.29 | 1.74 | 7 | 4.84 | 1.98 | 5 | 4.52 | 1.91 | 4 |
| Ordinary unfavorable | miner | -3.75 | 4.25 | 1.92 | 4 | 4.94 | 1.88 | 5 | 6 | 2.09 | 7 | 5.13 | 1.88 | 5 |
| Ordinary unfavorable | lazy employee | -7.06 | 3.09 | 2.26 | 2 | 2.73 | 2.44 | 2 | 6.7 | 2.26 | 7 | 6.45 | 1.92 | 6 |
| Ordinary unfavorable | sunburned person | -8.02 | 3.61 | 2.11 | 3 | 3.52 | 2.38 | 2 | 6.42 | 2.14 | 7 | 4.97 | 1.58 | 5 |
| Ordinary unfavorable | shooter | -4.06 | 2.66 | 2.34 | 1.5 | 5.19 | 2.18 | 5 | 7.69 | 1.42 | 8 | 3.25 | 2.18 | 3 |
| Ordinary unfavorable | armed robber | -5.46 | 2.33 | 2.06 | 2 | 5.39 | 2.5 | 6 | 8 | 1.71 | 9 | 3.24 | 2.18 | 3 |
| Ordinary unfavorable | landmine detector | -7.53 | 5.23 | 2.73 | 5 | 5.77 | 2.08 | 5 | 5.5 | 2.33 | 5 | 4.1 | 2.23 | 4 |
| Ordinary unfavorable | bandit | -4.03 | 3.19 | 2.3 | 3 | 5 | 2.22 | 5 | 6.35 | 2.26 | 7 | 4.32 | 2.02 | 4 |
| Ordinary favorable | butcher | -3.61 | 5.28 | 2.04 | 5 | 3.75 | 2.13 | 3 | 5.94 | 2.06 | 6 | 5.25 | 2.17 | 5 |
| Ordinary favorable | guitar player | -4.95 | 6.75 | 1.81 | 7 | 4.47 | 2.12 | 5 | 3.03 | 1.99 | 3 | 5.5 | 2.37 | 5 |
| Ordinary favorable | teacher | -2.16 | 6.78 | 1.88 | 7 | 3.59 | 2.43 | 3 | 3.5 | 1.93 | 3 | 5.84 | 2.14 | 6 |
| Ordinary favorable | skateboarder | -5.71 | 5.19 | 1.82 | 5 | 4.42 | 2.22 | 5 | 4.39 | 1.89 | 5 | 5.42 | 2.11 | 5 |
| Ordinary favorable | student | -2.08 | 6.66 | 2.01 | 7.5 | 2.5 | 1.83 | 2 | 3.44 | 2.2 | 3 | 5.72 | 2.53 | 5.5 |
| Ordinary favorable | public speaker | -4.64 | 5.88 | 2.03 | 6 | 3.97 | 2.07 | 4 | 4.59 | 1.97 | 4.5 | 4.94 | 2.15 | 5 |
| Ordinary favorable | crane operator | -5.36 | 4.74 | 1.84 | 5 | 4.26 | 2.24 | 4 | 5.23 | 1.91 | 5 | 5.87 | 2.01 | 6 |
| Ordinary favorable | working out | -3.56 | 7 | 2.05 | 8 | 3.03 | 1.87 | 3 | 3.52 | 2.41 | 3 | 4.39 | 1.89 | 5 |
| Ordinary favorable | businessman | -3.39 | 5.42 | 2.45 | 6 | 3.26 | 2.27 | 2 | 4.29 | 2.34 | 4 | 5.71 | 2.24 | 6 |
| Ordinary favorable | doctor | -2.36 | 7.19 | 2.02 | 8 | 4.91 | 2.66 | 5 | 3.16 | 2.03 | 2 | 5.28 | 2.34 | 5.5 |
| Ordinary favorable | ballet dancer | -4.79 | 6.58 | 1.58 | 6 | 5.42 | 2.18 | 6 | 3.55 | 2.02 | 3 | 5.88 | 2.25 | 6 |
| Ordinary favorable | shopper | -4.27 | 5.32 | 2.06 | 5 | 1.81 | 1.3 | 1 | 4.26 | 2.21 | 4 | 6.1 | 2.06 | 6 |
| Ordinary favorable | pilot | -2.77 | 6.23 | 1.94 | 6 | 5.48 | 2.28 | 6 | 3.71 | 1.9 | 4 | 4.94 | 2.32 | 5 |
| Ordinary favorable | fireman | -4.16 | 6.88 | 2.17 | 7.5 | 5.44 | 2.41 | 5.5 | 3.72 | 2.14 | 3 | 4.38 | 1.98 | 4 |
| Ordinary favorable | chef | -3.65 | 6.94 | 1.88 | 7 | 4.23 | 2.36 | 5 | 3.61 | 2.39 | 3 | 5.16 | 2.6 | 5 |
| Ordinary favorable | barista | -4.99 | 5.72 | 1.92 | 6 | 4.13 | 2.37 | 4 | 3.47 | 1.65 | 3.5 | 5.88 | 2.09 | 6 |
| Ordinary favorable | librarian | -3.42 | 6.5 | 1.76 | 7 | 3.27 | 2.23 | 3 | 3.5 | 1.63 | 3 | 7.13 | 1.63 | 7.5 |
| Ordinary favorable | gas pumper | -7.17 | 4.48 | 2.2 | 5 | 2.55 | 1.86 | 2 | 5.24 | 1.98 | 5 | 7.3 | 1.74 | 8 |
| Ordinary favorable | computer gamer | -7.4 | 4.71 | 2.15 | 5 | 3.32 | 1.94 | 3 | 5.35 | 2.04 | 5 | 5.35 | 2.12 | 5 |
| Ordinary favorable | astronaut | -4.08 | 7.18 | 1.72 | 8 | 7.12 | 1.58 | 8 | 2.85 | 1.52 | 2 | 4 | 2.06 | 4 |
| Ordinary favorable | gardener | -3.64 | 6.81 | 1.96 | 7 | 2.9 | 2.04 | 2 | 3.61 | 2.29 | 3 | 6.87 | 2.16 | 8 |
| Ordinary favorable | model | -1.69 | 5.56 | 1.97 | 5 | 4.78 | 2.38 | 5.5 | 3.88 | 2.11 | 4 | 5.56 | 2.42 | 5 |
| Ordinary favorable | chess player | -4.94 | 5.75 | 1.78 | 5.5 | 4.47 | 2.34 | 5 | 3.81 | 1.79 | 4 | 5.97 | 1.98 | 6 |
| Ordinary favorable | lab researcher | -7.11 | 6.33 | 1.97 | 7 | 4.73 | 1.96 | 5 | 4.47 | 2.06 | 5 | 5.83 | 1.91 | 6 |
| Ordinary favorable | hairdresser | -4.23 | 5.33 | 2.07 | 5 | 3.45 | 2.05 | 3 | 4.03 | 1.94 | 4 | 6.45 | 1.75 | 6 |
| Ordinary favorable | camper | -4.27 | 5.59 | 1.68 | 6 | 3 | 1.81 | 3 | 4.22 | 1.72 | 4.5 | 6.25 | 1.88 | 6.5 |
| Ordinary favorable | golfer | -4.14 | 5.55 | 1.68 | 5 | 3.39 | 2.16 | 3 | 4.64 | 1.98 | 5 | 6.61 | 1.54 | 7 |
| Ordinary favorable | bicyclist | -5.09 | 6.82 | 1.55 | 7 | 3.85 | 2.21 | 4 | 3.42 | 1.9 | 3 | 5.73 | 2.04 | 6 |
| Ordinary favorable | veterinarian | -3.92 | 6.72 | 1.84 | 7 | 4.69 | 1.77 | 5 | 3.59 | 1.7 | 3.5 | 5.75 | 1.65 | 5 |
| Ordinary favorable | dentist | -3.44 | 5.84 | 2.08 | 6 | 4.06 | 2.38 | 4 | 4.65 | 2.01 | 5 | 5.87 | 2 | 6 |
| Ordinary favorable | sushi chef | -6.14 | 6.3 | 1.93 | 6 | 4.82 | 2.26 | 5 | 3 | 1.58 | 3 | 5.36 | 2.32 | 5 |
| Ordinary favorable | hockey player | -5.01 | 5.87 | 2.01 | 6 | 5.03 | 2.46 | 5 | 4.1 | 2.26 | 4 | 4.7 | 2.32 | 5.5 |
| Ordinary favorable | photographer | -3.28 | 6.57 | 1.68 | 7 | 3.67 | 2.15 | 3.5 | 3.4 | 2.01 | 3 | 5.8 | 2.19 | 6.5 |
| Ordinary favorable | goal keeper | -6.28 | 6.55 | 1.66 | 6 | 4.28 | 2.33 | 5 | 4.66 | 2.24 | 5 | 5.9 | 1.76 | 5 |
| Ordinary favorable | yoga practitioner | -6.52 | 6.34 | 2.12 | 6 | 3.75 | 2.08 | 4 | 2.97 | 2.07 | 2 | 6.59 | 2.33 | 7 |
| Ordinary favorable | music listener | -6.39 | 7.16 | 1.92 | 7.5 | 2.75 | 2.74 | 1 | 2.34 | 1.66 | 2 | 6.53 | 2.4 | 7 |
| Ordinary favorable | toddler | -3.74 | 6.41 | 2.28 | 7 | 2.31 | 1.91 | 1 | 3 | 1.87 | 3 | 5.17 | 1.98 | 5 |
| Ordinary favorable | driving | -2.57 | 5.69 | 2.16 | 5 | 2.53 | 1.97 | 2 | 4.53 | 2.23 | 4.5 | 5.81 | 2.39 | 6 |
| Ordinary favorable | swimmer | -3.99 | 6.21 | 1.96 | 7 | 4.42 | 2 | 5 | 3.61 | 1.97 | 4 | 5.73 | 2.43 | 6 |
| Ordinary favorable | celebrator | -5.85 | 6.29 | 1.85 | 7 | 3.48 | 2.1 | 3 | 3.65 | 2.3 | 3 | 5.1 | 1.81 | 5 |
| Ordinary favorable | newly wed | -5.39 | 6.65 | 1.96 | 7 | 3.42 | 2.09 | 3 | 3.58 | 2.41 | 3 | 4.68 | 2.12 | 5 |
| Ordinary favorable | school child | -4.78 | 6.47 | 1.87 | 7 | 2.67 | 2.01 | 2 | 3.4 | 1.77 | 3 | 6.17 | 1.86 | 6 |
| Ordinary favorable | in love | -2.84 | 7.03 | 2.63 | 8 | 4.21 | 2.9 | 3 | 2.28 | 2.22 | 1 | 3.76 | 3 | 2 |
| Ordinary favorable | mother | -1.75 | 8.06 | 1.84 | 9 | 3.84 | 3.18 | 2 | 2.48 | 2.17 | 1 | 5.61 | 2.76 | 5 |
| Ordinary favorable | lawyer | -2.64 | 6 | 2.16 | 6 | 4.91 | 2.39 | 5 | 4.66 | 1.89 | 5 | 5.47 | 2.05 | 5 |
| Ordinary favorable | newspaper reader | -5.45 | 6.23 | 1.41 | 7 | 3.13 | 2.29 | 2 | 3.9 | 1.94 | 4 | 6.68 | 1.66 | 7 |
| Ordinary favorable | construction worker | -4.73 | 5.1 | 1.72 | 5 | 3.1 | 1.58 | 3 | 5.26 | 1.73 | 5 | 6.16 | 1.81 | 7 |
| Ordinary favorable | mountain climber | -5.15 | 6.28 | 2.04 | 6.5 | 5.72 | 2.45 | 7 | 3.44 | 1.72 | 3 | 4.25 | 2.27 | 4 |
| Ordinary favorable | judo wrestler | -7.5 | 5.61 | 1.91 | 5 | 5.58 | 2.05 | 6 | 4.65 | 1.72 | 5 | 4.9 | 2.62 | 5 |
| Ordinary favorable | baker | -3.89 | 6.62 | 1.9 | 7 | 3.17 | 2.14 | 2 | 2.83 | 1.47 | 3 | 6.38 | 2.29 | 7 |
| Ordinary favorable | grandmother | -2.94 | 7.25 | 2.17 | 8 | 3.38 | 2.71 | 2 | 2.44 | 1.61 | 2 | 5.97 | 2.43 | 5.5 |
| Ordinary favorable | grocery shopper | -6.53 | 5.59 | 1.79 | 5 | 2.53 | 2.33 | 1 | 4.19 | 1.77 | 4 | 6.28 | 2.19 | 6 |
| Ordinary favorable | computer engineer | -5.61 | 6.25 | 1.78 | 6 | 4.31 | 2.35 | 4 | 4.28 | 1.78 | 4.5 | 5.59 | 2.03 | 6 |
| Ordinary favorable | ice skating | -4.78 | 6.4 | 1.96 | 6.5 | 3.8 | 2.04 | 4 | 3 | 1.6 | 3 | 5.57 | 2.19 | 6 |
| Ordinary favorable | pregnant woman | -3.94 | 6 | 2.42 | 6 | 3.06 | 2.03 | 3 | 4.06 | 2.22 | 4 | 5 | 2.48 | 4 |
| Ordinary favorable | motorcyclist | -5 | 5.26 | 1.91 | 5 | 3.87 | 1.63 | 4 | 4.71 | 1.88 | 5 | 4.74 | 1.97 | 5 |
| Ordinary favorable | praying person | -6.11 | 5.44 | 2.29 | 5 | 3.31 | 2.04 | 2 | 4.03 | 2.36 | 4.5 | 6.28 | 2.59 | 7 |
| Ordinary favorable | professor | -2.73 | 6.45 | 1.8 | 7 | 5.03 | 2.2 | 5 | 4 | 1.67 | 4 | 5.03 | 1.92 | 5 |
| Ordinary favorable | football player | -4.35 | 5.56 | 2.61 | 6 | 4.44 | 2.2 | 5 | 4.63 | 2.55 | 4 | 5.34 | 2.12 | 5 |
| Ordinary favorable | botanist | -4.09 | 6.56 | 1.7 | 7 | 4.94 | 1.76 | 5 | 3.25 | 1.57 | 3 | 6.25 | 2.17 | 6.5 |
| Ordinary favorable | optometrist | -4.81 | 5.93 | 2.03 | 6 | 4.1 | 1.71 | 4 | 4.13 | 1.74 | 4.5 | 6.5 | 1.48 | 6 |
| Ordinary favorable | disc jockey | -4.7 | 5.6 | 2.25 | 6 | 4.57 | 2.22 | 5 | 4.43 | 2.53 | 5 | 4.43 | 2.08 | 4 |
| Ordinary favorable | singer | -3.17 | 6.53 | 2.16 | 7 | 5.16 | 1.92 | 5 | 3.56 | 1.83 | 4 | 5.19 | 2.25 | 5 |
| Ordinary favorable | cheerleader | -4.43 | 5.5 | 2.26 | 5.5 | 4.1 | 2.07 | 4 | 4.8 | 2.43 | 5 | 5.03 | 2.14 | 5 |
| Ordinary favorable | weightlifter | -5.42 | 5.38 | 1.93 | 5 | 5.14 | 1.98 | 6 | 5.03 | 1.68 | 5 | 5.28 | 2.09 | 5 |
| Ordinary favorable | sailor | -3.45 | 6.13 | 1.43 | 6 | 5.06 | 1.65 | 5 | 3.68 | 1.51 | 4 | 5.52 | 1.93 | 5 |
| Ordinary favorable | bartender | -3.88 | 5.75 | 2.05 | 6 | 3.94 | 2.18 | 4 | 4.09 | 2.1 | 4 | 5.53 | 1.8 | 5 |
| Ordinary favorable | police officer | -3.6 | 5.9 | 2.18 | 7 | 3.9 | 2.21 | 3 | 5.23 | 2.33 | 5 | 4.55 | 2.14 | 5 |
| Ordinary favorable | soldier | -2.78 | 4.9 | 2.4 | 5 | 5.19 | 1.96 | 5 | 5.84 | 2.66 | 6 | 4.13 | 2.25 | 4 |
| Ordinary favorable | fisherman | -3.62 | 5.88 | 2.15 | 6 | 3.85 | 2.36 | 3 | 4.52 | 2.05 | 4 | 6.21 | 1.93 | 6 |
| Ordinary favorable | trail biker | -7.73 | 5.69 | 1.82 | 6 | 4.47 | 1.72 | 5 | 4.31 | 2.07 | 4 | 5.16 | 1.92 | 5 |
| Ordinary favorable | canoeist | -5.33 | 6.12 | 1.36 | 5 | 4.24 | 1.52 | 4 | 3.91 | 1.44 | 4 | 6.3 | 1.72 | 6 |
| Ordinary favorable | scientist | -2.99 | 7.03 | 1.8 | 7 | 5.59 | 2.17 | 6.5 | 4.16 | 2.13 | 4.5 | 5.06 | 1.92 | 5 |
| Ordinary favorable | marathon runner | -5.25 | 6.36 | 1.67 | 6 | 5.42 | 2.25 | 5 | 4.09 | 1.77 | 4 | 5 | 2.25 | 5 |
| Ordinary favorable | tennis player | -4.6 | 6.58 | 1.82 | 6 | 4.58 | 2.17 | 5 | 3.94 | 1.98 | 4 | 5.29 | 1.99 | 5 |
| Ordinary favorable | horse rider | -5.93 | 6.16 | 1.69 | 6 | 4.22 | 2.09 | 4.5 | 3.59 | 1.93 | 3.5 | 5.09 | 1.8 | 5 |
| Ordinary favorable | flight hostess | -7.32 | 6.42 | 1.78 | 7 | 3.32 | 2.1 | 3 | 4.35 | 2.33 | 4 | 6.23 | 1.73 | 6 |
| Ordinary favorable | politician | -3.18 | 5 | 1.86 | 5 | 4.35 | 2.51 | 4 | 5.48 | 2.06 | 5 | 4.74 | 1.88 | 5 |
| Ordinary favorable | baseball player | -4.49 | 5.3 | 2 | 5 | 4.5 | 2.33 | 4.5 | 4.83 | 2.05 | 4 | 6.3 | 1.97 | 6 |
| Ordinary favorable | gymnast | -4.54 | 6.26 | 1.65 | 6 | 5.55 | 2.22 | 6 | 3.39 | 1.82 | 4 | 5 | 1.79 | 5 |
| Ordinary favorable | newborn baby | -4.53 | 7.13 | 1.93 | 8 | 3.6 | 2.79 | 2 | 3.2 | 2.4 | 2 | 5.47 | 2.29 | 5 |
| Ordinary favorable | father | -1.78 | 7.78 | 1.86 | 9 | 3.13 | 2.93 | 1.5 | 2.66 | 1.98 | 2 | 5.97 | 2.12 | 6 |
| Ordinary favorable | dog owner | -5.2 | 6.03 | 1.94 | 6 | 2.5 | 1.83 | 2 | 3.43 | 1.96 | 3 | 5.97 | 1.77 | 6 |
| Ordinary favorable | waterpolo player | -7.94 | 5.53 | 1.74 | 5 | 4.2 | 1.81 | 5 | 3.83 | 1.62 | 4 | 5.53 | 1.94 | 5 |
| Ordinary favorable | lacrosse player | -6.15 | 5.38 | 2 | 5 | 4.41 | 1.74 | 5 | 4.78 | 2.12 | 5 | 4.97 | 2.15 | 5 |
| Ordinary favorable | painter | -3.01 | 6.27 | 1.31 | 6 | 3.6 | 2.24 | 4 | 3.77 | 1.81 | 3.5 | 6.2 | 1.86 | 6 |
| Ordinary favorable | secretary | -2.62 | 5.45 | 1.55 | 5 | 2.81 | 2.1 | 2 | 4.58 | 1.8 | 5 | 6.84 | 1.77 | 7 |
| Ordinary favorable | trumpeter | -4.38 | 5.7 | 1.95 | 6 | 4.37 | 2.16 | 4 | 4 | 2.07 | 4 | 5.97 | 1.73 | 6 |
| Ordinary favorable | club dancer | -6.72 | 4.88 | 2.15 | 5 | 3.21 | 2.1 | 3 | 4.67 | 2.19 | 4 | 4.7 | 2.39 | 5 |
| Ordinary favorable | security guard | -4.25 | 5.35 | 2.03 | 5 | 3.23 | 1.82 | 3 | 5.16 | 2.22 | 5 | 5.71 | 2.37 | 6 |
| Ordinary favorable | skier | -4.29 | 5.67 | 1.69 | 5 | 4.03 | 1.79 | 5 | 4.24 | 1.8 | 4 | 5.48 | 1.8 | 5 |
| Ordinary favorable | rower | -5 | 5.97 | 1.69 | 6 | 4.24 | 2.08 | 4 | 4.15 | 1.62 | 4 | 5.7 | 2.02 | 5 |
| Ordinary favorable | boxer | -4.01 | 5 | 1.57 | 5 | 5.03 | 2.15 | 5 | 5.32 | 1.87 | 5 | 4.39 | 1.99 | 5 |
| Ordinary favorable | bowler | -4.29 | 5.47 | 1.68 | 5 | 3.44 | 2.03 | 3 | 4.22 | 1.93 | 4 | 6.38 | 2.04 | 6.5 |
| Ordinary favorable | farmer | -2.85 | 6.31 | 2.12 | 7 | 3.84 | 1.87 | 4 | 4.19 | 1.94 | 4 | 6.09 | 1.94 | 6 |
| Ordinary favorable | violinist | -4 | 6.53 | 1.57 | 6.5 | 4.8 | 2.33 | 5 | 3.03 | 1.69 | 3 | 6.27 | 2.16 | 7 |
| Ordinary favorable | basketball player | -4.56 | 5.26 | 2.25 | 5 | 4.94 | 2.29 | 5 | 3.97 | 2.18 | 4 | 5.23 | 1.96 | 5 |
| Ordinary favorable | pole vaulter | -5.88 | 6 | 1.81 | 6 | 6.5 | 1.74 | 7 | 4.44 | 1.95 | 4 | 5.53 | 2.29 | 5.5 |
| Ordinary favorable | parachutist | -5.04 | 5.82 | 1.94 | 6 | 6.33 | 1.85 | 7 | 4.61 | 1.95 | 5 | 4 | 2.29 | 4 |
| Ordinary favorable | motorcycle racer | -6.35 | 5.1 | 2.26 | 5 | 5.26 | 2.16 | 6 | 4.42 | 2.14 | 4 | 4.23 | 1.89 | 4 |
| Ordinary favorable | lover | -2.87 | 7.1 | 2.33 | 8 | 4.16 | 2.86 | 3 | 2.03 | 1.52 | 1 | 3.58 | 2.6 | 3 |
| Ordinary favorable | curler | -5.49 | 5.36 | 1.65 | 5 | 4 | 1.71 | 4 | 4.27 | 1.72 | 5 | 6.48 | 1.82 | 7 |
| Ordinary favorable | jet skier | -7.34 | 5.91 | 1.96 | 6 | 4.67 | 1.95 | 5 | 3.73 | 1.82 | 3 | 4.7 | 2.13 | 5 |
| Ordinary favorable | judge | -2.38 | 5.94 | 2.01 | 6 | 4.76 | 2.22 | 5 | 5.18 | 1.79 | 5 | 5.36 | 1.9 | 5 |
| Ordinary favorable | diver | -3.87 | 6.23 | 1.5 | 6 | 5.16 | 1.95 | 5 | 4.26 | 1.88 | 4 | 4.94 | 2.03 | 5 |
| Ordinary favorable | surfer | -4.5 | 6.19 | 1.97 | 7 | 4.29 | 2 | 5 | 3.74 | 1.86 | 3 | 4.77 | 2.06 | 5 |
| Ordinary favorable | trophy winner | -6.66 | 5.83 | 2.39 | 6.5 | 5.07 | 2.27 | 5 | 4.07 | 2.24 | 4 | 4.53 | 1.94 | 4.5 |
| Ordinary favorable | rafter | -4.32 | 5.53 | 1.5 | 5 | 4.56 | 1.87 | 5 | 4.28 | 1.69 | 5 | 5.06 | 2.18 | 5 |
| Ordinary favorable | birthday girl | -5.64 | 6.42 | 2.23 | 7 | 2.9 | 2.52 | 2 | 2.9 | 2.13 | 2 | 5.39 | 2.54 | 5 |
| Ordinary favorable | rock climber | -5.61 | 5.9 | 1.68 | 6 | 5.19 | 2.17 | 5 | 3.77 | 1.86 | 4 | 4.1 | 2.07 | 4 |
| Ordinary favorable | zookeeper | -5.47 | 5.87 | 2.11 | 6 | 4.9 | 2.13 | 5 | 3.84 | 1.71 | 4 | 5.61 | 1.8 | 6 |
| Ordinary favorable | sports fan | -5.25 | 5.13 | 2.23 | 5 | 2.68 | 2.1 | 2 | 4.39 | 2.08 | 4 | 5.35 | 2.07 | 5 |
| Ordinary favorable | pharmacist | -3.81 | 6.1 | 1.73 | 6 | 3.63 | 2.04 | 3 | 4 | 1.64 | 4 | 6.13 | 1.96 | 6.5 |
| Ordinary favorable | concert goer | -6.86 | 5.94 | 1.61 | 5 | 3.29 | 1.9 | 3 | 3.71 | 1.85 | 4 | 5.23 | 2.01 | 5 |
| Ordinary favorable | massage client | -7.3 | 5.81 | 1.94 | 6 | 3.55 | 1.96 | 3 | 3.84 | 2.11 | 4 | 6.32 | 2.27 | 7 |
| Ordinary favorable | movie goer | -6.83 | 6.41 | 1.98 | 7 | 2.03 | 1.62 | 1 | 3.34 | 1.84 | 3 | 7.13 | 1.64 | 8 |
| Ordinary favorable | carpenter | -3.54 | 5.88 | 1.73 | 6 | 4 | 2.06 | 4 | 4.12 | 1.24 | 4 | 6.09 | 1.96 | 6 |
| Ordinary favorable | restaurant customer | -6.58 | 5.52 | 1.91 | 5 | 2.45 | 1.91 | 2 | 4.13 | 2.03 | 5 | 6.23 | 1.96 | 6 |
| Ordinary favorable | dancer | -3.4 | 6.52 | 1.92 | 7 | 4.94 | 2.42 | 5 | 3.58 | 2.14 | 3 | 4.79 | 2.1 | 4 |
| Ordinary favorable | nature lover | -5.55 | 6.9 | 1.99 | 8 | 4.03 | 2.07 | 4 | 3.19 | 1.96 | 3 | 6 | 2.05 | 6 |
| Ordinary favorable | piano teacher | -4.96 | 6.34 | 1.99 | 6.5 | 4.25 | 2.02 | 4.5 | 3.56 | 1.76 | 3 | 6.63 | 1.6 | 7 |
| Ordinary favorable | jogger | -4.76 | 5.77 | 2.09 | 6 | 3.29 | 1.77 | 3 | 3.97 | 1.85 | 4 | 5.87 | 1.88 | 6 |
| Ordinary favorable | sun tanning | -6.38 | 4.03 | 2.67 | 3 | 2.63 | 2.12 | 2 | 5.28 | 2.47 | 5 | 5.78 | 2.2 | 6 |
| Ordinary favorable | piano student | -5.86 | 6.48 | 1.73 | 6 | 4.29 | 1.95 | 5 | 3.84 | 1.92 | 4 | 6.84 | 1.92 | 7 |
| Ordinary favorable | snowboarder | -5.79 | 6.16 | 1.7 | 6 | 4.29 | 2.24 | 5 | 4.42 | 2.32 | 5 | 5.39 | 1.78 | 5 |
| Ordinary favorable | texter | -6.28 | 5.16 | 1.98 | 5 | 2.48 | 2.5 | 1 | 4.87 | 1.86 | 5 | 6.29 | 2.3 | 7 |
| Ordinary favorable | roller skater | -6.39 | 5.73 | 1.53 | 5 | 3.73 | 1.99 | 4 | 4.24 | 1.7 | 4 | 5.55 | 1.82 | 5 |
| Ordinary favorable | tree planter | -6.32 | 6.63 | 2.24 | 7 | 4.16 | 2.02 | 4 | 3.28 | 2.07 | 3 | 5.75 | 2.26 | 6 |
| Ordinary favorable | mechanic | -3.61 | 5.61 | 2 | 6 | 3.64 | 1.75 | 4 | 4.76 | 1.75 | 5 | 5.85 | 1.72 | 6 |
| Ordinary favorable | meditator | -3.43 | 6.63 | 1.54 | 7 | 4.38 | 2.11 | 5 | 3.06 | 1.85 | 2 | 6.56 | 2.14 | 6.5 |
| Ordinary favorable | pub customer | -7.69 | 5.62 | 1.57 | 6 | 2.83 | 1.75 | 3 | 4.72 | 1.53 | 5 | 5.69 | 1.63 | 6 |
| Ordinary favorable | celebrating Christmas | -5.56 | 7.03 | 2.21 | 8 | 3.19 | 2.48 | 2 | 2.94 | 2.57 | 2 | 4.26 | 2.68 | 3 |
| Ordinary favorable | office worker | -4.81 | 5.34 | 2.46 | 5.5 | 2.56 | 2.02 | 2 | 5.25 | 2.05 | 5 | 6.78 | 1.72 | 7 |
| Ordinary favorable | pool player | -5.81 | 5.52 | 2.29 | 5 | 3.26 | 1.73 | 3 | 4.06 | 2.02 | 4 | 6.32 | 1.83 | 7 |
| Ordinary favorable | dog walker | -6 | 6.19 | 1.87 | 6 | 2.84 | 2.07 | 2 | 3.45 | 1.48 | 3 | 6.87 | 1.59 | 7 |
| Ordinary favorable | sprinter | -4.77 | 6.19 | 2.02 | 6 | 5.66 | 2.09 | 6 | 4.03 | 1.84 | 4 | 5.28 | 2.11 | 5 |
| Ordinary favorable | surgeon | -3.04 | 6.59 | 1.84 | 7 | 5.38 | 2.58 | 6 | 3.9 | 2.16 | 3 | 5.31 | 2.19 | 5 |
| Ordinary favorable | jeweler | -4.37 | 5.84 | 1.59 | 6 | 3.88 | 1.79 | 4 | 3.97 | 1.49 | 4 | 6.16 | 1.97 | 6.5 |
| Ordinary favorable | tourist | -3.04 | 5.53 | 2.18 | 5.5 | 3.56 | 2.29 | 3 | 4.47 | 2.12 | 5 | 5.66 | 1.89 | 6 |
| Ordinary favorable | clown | -3.72 | 4.26 | 2.19 | 4 | 5.35 | 1.92 | 5 | 5.9 | 2.23 | 6 | 5.23 | 2.11 | 5 |
| Ordinary favorable | shepherd | -3.46 | 5.41 | 1.9 | 5 | 4.31 | 1.97 | 5 | 4.16 | 1.94 | 4 | 6.16 | 2.03 | 6 |
| Ordinary favorable | graduate | -2.76 | 6.87 | 1.98 | 7 | 3.84 | 2.11 | 4 | 3.26 | 1.75 | 3 | 5.39 | 2.28 | 5 |
| Ordinary favorable | stockholder | -3.85 | 5.47 | 2 | 5 | 3.38 | 1.77 | 3 | 4.88 | 1.79 | 5 | 5.69 | 2.05 | 5 |
| Ordinary favorable | detective | -3.22 | 6.56 | 1.54 | 7 | 5.53 | 1.83 | 6 | 4.06 | 1.81 | 3 | 4.72 | 1.63 | 5 |
| Ordinary favorable | lawn mower | -4.66 | 4.94 | 2.3 | 5 | 3.06 | 2.34 | 2 | 5.28 | 2.17 | 5 | 6.69 | 1.94 | 7 |
| Ordinary favorable | pilgrim | -3.77 | 5.73 | 1.48 | 6 | 5.2 | 2.12 | 6 | 4.67 | 2.14 | 5 | 6.13 | 2.08 | 6 |
| Ordinary favorable | archer | -4.36 | 5.53 | 1.83 | 5 | 5.33 | 1.58 | 5.5 | 4 | 1.72 | 4 | 5.07 | 1.93 | 5 |
| Ordinary favorable | paramedic | -4.56 | 6.61 | 2.36 | 7 | 4.94 | 2.14 | 5 | 3.48 | 2.2 | 3 | 4.61 | 2.12 | 4 |
| Ordinary favorable | polo player | -4.48 | 5.81 | 2.01 | 6 | 4.63 | 2.31 | 5 | 4.41 | 2.01 | 4 | 6.28 | 2.3 | 7 |
| Ordinary favorable | TV watcher | -6.42 | 4.84 | 1.83 | 5 | 2.65 | 2.47 | 1 | 4.23 | 2 | 5 | 6.58 | 2.36 | 7 |
| Ordinary favorable | architect | -3 | 6.68 | 1.68 | 7 | 4.87 | 2.28 | 5 | 3.06 | 1.31 | 3 | 5.87 | 1.8 | 6 |
| Ordinary favorable | forest ranger | -5.27 | 6.22 | 1.6 | 6 | 4.94 | 1.95 | 5 | 3.78 | 1.9 | 3 | 5.13 | 1.81 | 5 |
| Ordinary favorable | cowboy | -3.52 | 5.8 | 1.99 | 6 | 4.57 | 2.27 | 5 | 4.23 | 2.06 | 4 | 5.1 | 1.92 | 5 |
| Ordinary favorable | personal trainer | -5.15 | 6.1 | 2.44 | 6.5 | 3.87 | 1.93 | 4 | 4.2 | 2.06 | 4 | 5.4 | 1.89 | 5 |
| Ordinary favorable | journalist | -3.13 | 6.26 | 2.13 | 7 | 4.65 | 2.24 | 5 | 3.68 | 1.99 | 3 | 5.26 | 1.97 | 5 |
| Ordinary favorable | hiker | -4.53 | 6 | 1.44 | 6 | 3.66 | 1.77 | 4 | 3.94 | 1.61 | 4.5 | 5.91 | 1.8 | 6 |
| Ordinary favorable | crossing guard | -5.57 | 5.94 | 2.21 | 6 | 3.42 | 1.91 | 4 | 4.26 | 2.28 | 4 | 6.35 | 1.68 | 7 |
| Ordinary favorable | butterfly catcher | -7.06 | 5.61 | 2.38 | 6 | 5.16 | 2.46 | 5 | 3.84 | 2.38 | 3 | 5.97 | 2.12 | 6 |
| Ordinary favorable | presenter | -4.17 | 6 | 1.73 | 6 | 3.71 | 2.08 | 3 | 4.39 | 2.25 | 5 | 6.29 | 1.99 | 7 |
| Ordinary favorable | chemist | -3.63 | 6.06 | 1.95 | 7 | 4.63 | 2.12 | 5 | 4.41 | 1.93 | 4 | 5.72 | 1.63 | 5 |
| Ordinary favorable | radio show host | -6.49 | 6.03 | 2.15 | 7 | 4.32 | 2.07 | 5 | 3.71 | 2.24 | 3 | 5.58 | 1.84 | 5 |
| Ordinary favorable | nun | -3.49 | 4.97 | 2.61 | 5 | 4.61 | 2.08 | 5 | 4.77 | 2.59 | 5 | 6.61 | 2.11 | 7 |
| Ordinary favorable | priest | -2.68 | 4.75 | 2.41 | 5 | 5.03 | 2.42 | 5 | 5.25 | 2.74 | 6 | 5.97 | 2.31 | 5.5 |
| Ordinary favorable | social worker | -3.49 | 6.48 | 1.95 | 7 | 4.61 | 1.89 | 5 | 3.42 | 2.17 | 3 | 5.9 | 1.8 | 6 |
| Ordinary favorable | astronomer | -3.81 | 6.75 | 1.9 | 7.5 | 5.75 | 2.34 | 7 | 3.44 | 1.81 | 3 | 5.28 | 2.22 | 5 |
| Ordinary favorable | aircraft controller | -6.99 | 6.63 | 1.88 | 7 | 5.19 | 2.35 | 6 | 3.88 | 1.72 | 4 | 5.69 | 2.15 | 6 |
| Ordinary favorable | archeologist | -4.73 | 6.32 | 2.01 | 7 | 5.16 | 2.31 | 5 | 3.39 | 1.73 | 4 | 5.23 | 2.12 | 5 |
| Ordinary favorable | bus driver | -4.38 | 5.48 | 1.89 | 6 | 3.33 | 2.04 | 3 | 4.79 | 1.76 | 5 | 7 | 1.25 | 7 |
| Ordinary favorable | train conductor | -5.62 | 5.56 | 1.37 | 6 | 3.41 | 1.78 | 3 | 4.59 | 1.66 | 4 | 6.22 | 1.43 | 6 |
| Ordinary favorable | wine taster | -6.09 | 6.22 | 1.62 | 6 | 4.31 | 2.29 | 5 | 3.75 | 1.9 | 3.5 | 6.34 | 1.91 | 7 |
| Ordinary favorable | manicurist | -5.23 | 5.23 | 2.26 | 5 | 3.65 | 1.92 | 4 | 4.81 | 2.3 | 5 | 6.52 | 1.93 | 6 |
| Ordinary favorable | Nobel Peace Prize recipient | -6.5 | 7.68 | 2.07 | 9 | 6.83 | 2.38 | 8 | 2.32 | 2.09 | 1 | 4.32 | 2.29 | 4 |
| Ordinary favorable | movie director | -5.2 | 6.84 | 1.42 | 7 | 5.84 | 2.16 | 7 | 3.55 | 1.89 | 3 | 4.39 | 1.93 | 4 |
| Ordinary favorable | traveler | -3.52 | 6.94 | 1.73 | 7 | 4.16 | 2.42 | 5 | 3.13 | 2.25 | 3 | 4.58 | 2.62 | 4 |
| Ordinary favorable | story teller | -5.1 | 6.88 | 1.72 | 7 | 4 | 2.09 | 4 | 3.03 | 1.6 | 2.5 | 6.13 | 2.25 | 6.5 |
| Ordinary favorable | nurse | -2.6 | 7.16 | 1.87 | 7 | 4.03 | 2.26 | 4 | 3.34 | 1.84 | 3 | 5.16 | 1.87 | 5 |
| Ordinary favorable | voter | -3.41 | 6.74 | 2.13 | 8 | 3 | 2.07 | 2 | 3.97 | 2.21 | 3 | 5.84 | 2.24 | 6 |
| Ordinary favorable | writer | -2.36 | 6.87 | 2.16 | 7 | 4.77 | 2.17 | 5 | 3.32 | 1.8 | 3 | 5.52 | 2.25 | 5 |
| Ordinary favorable | playing child | -6.06 | 7.4 | 1.4 | 8 | 2.7 | 2.37 | 2 | 2.5 | 2 | 2 | 5.87 | 2.21 | 6 |
| Ordinary favorable | lifeguard | -4.56 | 7.19 | 1.53 | 8 | 4.38 | 2.15 | 4 | 3.16 | 1.67 | 3 | 5.69 | 1.97 | 5.5 |
| Ordinary favorable | griller | -6.04 | 5.3 | 1.97 | 5 | 3.57 | 2.31 | 3 | 4.33 | 1.9 | 4 | 6.43 | 1.48 | 6 |
| Ordinary favorable | body surfer | -6.77 | 5.29 | 1.37 | 5 | 4.71 | 2.04 | 5 | 4.48 | 1.48 | 5 | 5.42 | 1.77 | 5 |
| Ordinary favorable | blogger | -5.38 | 5.16 | 2.1 | 5 | 3.19 | 2.07 | 3 | 4.29 | 1.79 | 5 | 6.48 | 1.55 | 6 |
| Ordinary favorable | playing paintball | -7.11 | 5.13 | 2.2 | 5 | 3.71 | 2.22 | 4 | 5.06 | 2.19 | 6 | 3.94 | 2.17 | 3 |
| Ordinary favorable | dinner host | -6.56 | 6.06 | 2.18 | 6 | 2.94 | 2.02 | 2.5 | 3.47 | 1.9 | 3 | 5.78 | 1.75 | 5 |
| Ordinary favorable | X-ray technician | -5.94 | 5.63 | 1.85 | 6 | 4.4 | 2.09 | 5 | 4.13 | 1.33 | 5 | 6.03 | 1.94 | 6.5 |
